# Chloroplast- and Mitochondrion-Specific Random C-to-T Mutagenesis for Forward Genetics of Organelle Genomes

**DOI:** 10.1101/2025.11.01.685978

**Authors:** Nanami Kosaka, Yoshiki Harada, Issei Nakazato, Miki Okuno, Takehiko Itoh, Nobuhiro Tsutsumi, Shin-ichi Arimura

## Abstract

Organelle genomes in plastids (including chloroplasts) and mitochondria encode essential genes for photosynthesis, respiration, and agronomic traits, representing promising targets for crop improvement. However, their high copy number and non-Mendelian inheritance have long hindered efficient modification compared to nuclear genomes. Recent advances in organelle base editing (C-to-T and A-to-G) have enabled precise nucleotide substitutions, yet information on useful mutations remains limited. Here, to establish a forward genetics platform for C-to-T substitutions, we developed a method to introduce random C-to-T mutations throughout the entire organelle genomes of *Arabidopsis thaliana*. We engineered a fusion protein, WHY2-CD mutator, combining cytidine deaminase (CD), uracil glycosylase inhibitor, and sequence-nonspecific DNA-binding protein WHIRLY2 (WHY2), fused to organelle-targeting peptides. This system introduced dispersed C-to-T substitutions specifically within plastid or mitochondrial genomes. In T_2_ lines, we identified homoplasmic (homozygous) plastid genome mutants, including *rpoA* knockouts and *rbcL* variants with an amino acid substitution. Screening T_3_ populations on spectinomycin revealed plastid genome mutants with resistant traits and their causal mutation. These mutations can be transferred or combined using targeted base editors, such as transcription activator-like effector cytidine deaminase (TALECD). This comprehensive, C-to-T-focused mutagenesis provides a powerful tool for organelle forward genetics and molecular breeding.

## Introduction

The plastid genome of flowering plants encodes genes involved in photosynthesis, while the mitochondrial genome harbors genes for respiratory chain complexes, cytoplasmic male sterility (CMS), and other essential functions^1–3^. Variation in these organelle genomes and their interactions with the nuclear genome can influence field phenotypes^4–7^, and subtle or single nucleotide polymorphisms within organelle genomes have been linked to traits such as CMS^8^ and herbicide resistance^9, 10^. Thus, the ability to modify and precisely control organelle genomes holds great potential for improving crop performance; therefore, such methods could be a promising tool for crop breeding. Nevertheless, the use of organelle genomes has been limited in breeding, except for exploiting CMS in F_1_ hybrid seed production. Because organelle genomes are typically inherited uniparentally, creating a new combination of genes between the parental alleles and trait mapping is not feasible through crossing. Moreover, their high copy number per cell can make it difficult to achieve homoplasmy of introduced alleles or mutations, further complicating genetic analysis and breeding.

In recent years, plant organelle genome editing has advanced rapidly^3, 11^, enabling both targeted DNA cleavage-based editing^12^ and targeted base substitution^13–17^. These technologies have been applied to identify CMS-associated genes in the mitochondrial genome of rice & rapeseed^12^, tomato^18^, broccoli^19^, potato^20^, and eggplant^21^, to confer resistance to herbicides such as metribuzin and atrazine in Arabidopsis^22, 23^, and to improve photosynthetic capacity in Arabidopsis^24^. Currently, organelle-targeted base editing is largely limited to C-to-T and A-to-G substitutions, but due to the minimal alterations and the fact that such edits seem more acceptable to society and are often exempt from GMO regulations in some countries, this technology holds great promise for improving crop productivity. While the sequence information of organelle genomes is abundant^3^, data on naturally occurring or targetable base polymorphisms—particularly those that lead to phenotypic changes upon base editing—remain critically scarce^25–27^. This lack of information currently represents a major bottleneck for the effective application of genome editing in crop trait improvement.

Forward genetics by irradiation and chemical mutagenesis has long served as a powerful tool in the nucleus; numerous nuclear genome mutants have been generated and analyzed for identifying gene functions of interest and enhancing crop performance via mutational breeding. In contrast, reports on organelle genome mutants with a point mutation remain extremely limited, with only a few examples, such as *rbcL* mutants deficient in Rubisco holoenzyme in *Nicotiana tabacum* induced by ethyl methane sulfonate (EMS)^28^, lincomycin-resistant mutants in *Capsium annuum* by EMS^29^, and pigment-deficient or spectinomycin-resistant mutants by nitroso-methyl urea (NMU)^30–33^. Ethidium bromide increased the appearance of mitochondrial *nad9* point mutations under gene-drive-like circumstance^34^. Since chemical mutagens introduce mutations simultaneously in the nucleus, efficient isolation of organelle genome mutants is technically challenging. Some nuclear genome mutants are known to impair organelle DNA repair pathways, leading to the accumulation of mutations in organelle genomes. A representative example is the *MutS Homolog 1* (*MSH1*) mutant, originally isolated as *chloroplast mutator* (*chm*)^35–39^, which exhibits elevated point mutation frequencies in both plastid and mitochondrial genomes^40, 41^. However, *msh1* mutants induce mutations simultaneously in both organelle genomes, and each mutation is often not able to be transplanted separately to other plants. Furthermore, *msh1* can also trigger extensive genomic rearrangements via illegitimate homologous recombination between repeated sequences in the mitochondrial genome^42^. Attempts have also been made to generate a plastid genome mutant library using transposon tagging in tobacco, but the insertions tend to occur at limited specific loci with high specificity^43^. TALE nuclease (TALEN) gene-drive mutagenesis^34^ introduces mutations but it is limited to the TALE binding site.

In this study, we propose a method to introduce random C-to-T mutagenesis into the entire organelle genomes of *Arabidopsis thaliana*, for their forward genetic analysis, which can be applied to mutational breeding of crop organelle genomes. We employed a construct termed “WHY2-CD mutator,” which was made by fusing CD, UGI, a sequence-nonspecific DNA-binding protein AtWHIRLY2 (WHY2)^44, 45^, and a transit peptide for plastids or mitochondria^13, 46^, to induce C-to-T base substitutions at undefined loci in the targeted organelle genomes. Since double-stranded DNA deaminase toxin A (DddA) was adopted as CD^47^, due to its intrinsic preference for inducing tC-to-tT substitutions, the “TC” context is a potential editing site. Although the sites of mutagenesis are limited to specific contexts, the resulting variants are expected to be transferable or re-introducible into wild-type or other crop species using precise C-to-T base-editing by ptp/mitoTALECD^13, 15^. Here, we attempted to introduce transferable ‘semi’-random C-to-T mutations in a plastid- or mitochondrion-specific manner via signal peptide targeting, and to isolate mutants from a population of plastid genome-mutagenized lines.

## Results

### Construction of organelle-specific C-to-T mutagen vectors

To induce mutations in organelle genomes, binary vectors for *Agrobacterium*-mediated transformation were constructed to express artificially designed proteins, termed “WHY2-CD mutators.” These proteins consist of a mitochondrial or plastid targeting peptide, *Arabidopsis thaliana* Whirly2, which binds DNA without sequence specificity^45^, and the double-stranded DNA deaminase toxin A (DddA) from *Burkholderia cenocepacia*^47^ (Fig. 1A). Attempts to clone WHY2-CD mutators containing the wild-type, full-length DddA were unsuccessful (Supplementary Fig. S1), likely due to the high cytotoxicity and excessive substitution activity of DddA^47^. To attenuate the base substitution activity of DddA in the expression constructs, we employed two strategies: (1) splitting DddA at the 1,397th amino acid residue^47^, and (2) using less active DddA variants, E1347D and GSVG^48^ (Supplementary Fig. S2). Ultimately, eight WHY2-CD mutator constructs with different configurations were successfully designed, cloned, and introduced into plants (Fig. 1B). DddA prefers 5’-TC-3’ sequence contexts for cytosine substitution. In *Arabidopsis thaliana* ecotype Columbia-0 (Col-0), 52,381 and 20,219 “TC” sites are contained in the mitochondrial and plastid genomes, respectively (Fig. 1C), representing potential targets for the mutator. Of these, 5,951 and 7,587 sites in the mitochondrial and plastid genomes, respectively, are located within coding regions or open reading frames (ORFs) and may result in amino acid substitutions (Fig. 1D).

**Fig. 1.**
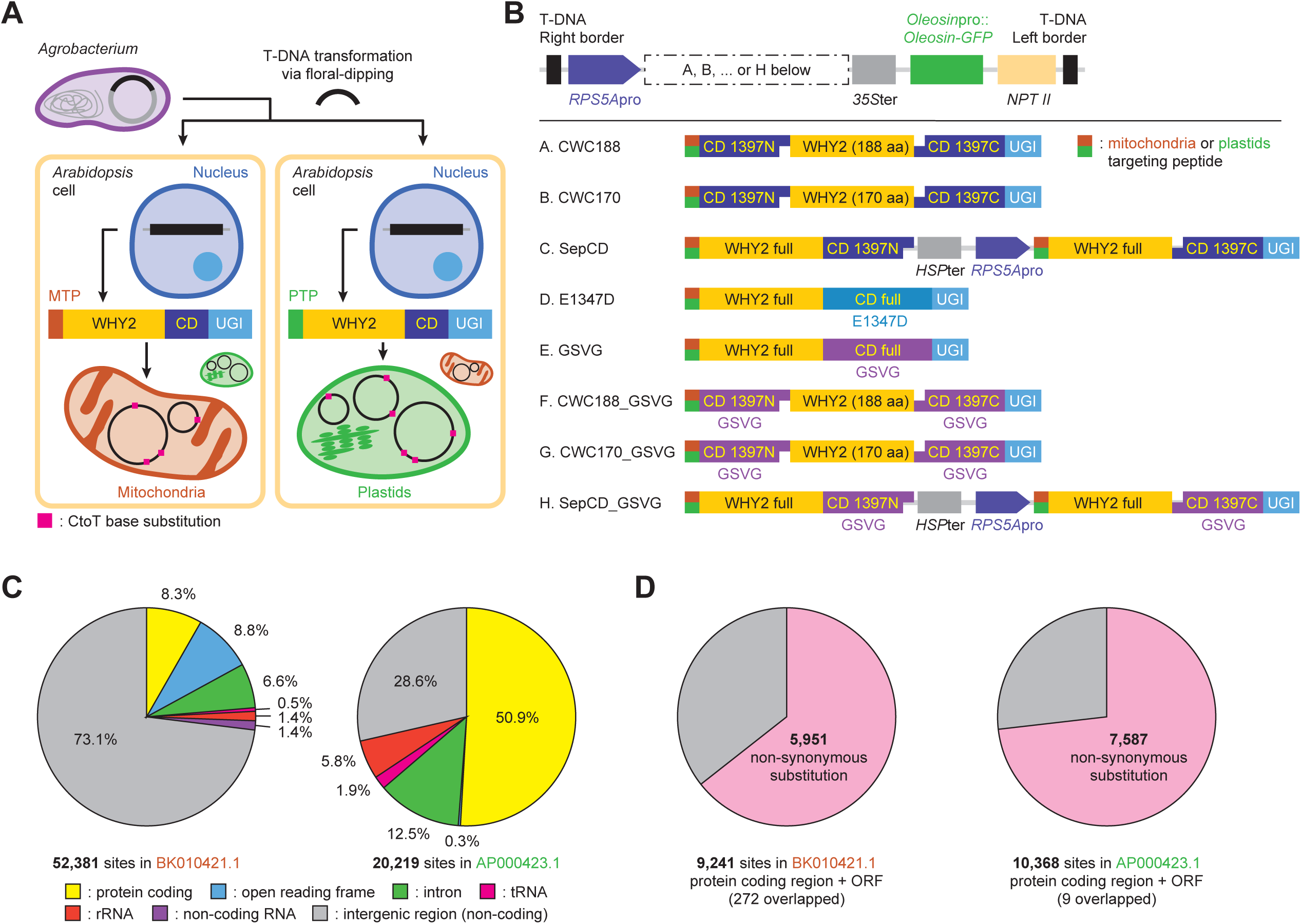
Organelle random mutagenesis strategy and scope of potential targets to be mutated. **A.** A schematic illustration of “WHY2-CD mutator” strategy. MTP, mitochondria targeting peptide; PTP, plastids targeting peptide; WHY2, Whirly2; CD, cytidine deaminase; UGI, uracil glycosylase inhibitor. **B.** Gene structures of each mutator construct. pro, promoter; ter, terminator; NPT, neomycin phosphor-transferase. For amino acid sequences of shorter WHY2, see Supplementary Information. **C.** The number of “TC” contexts and the proportion associated with each genetic element on the *A. thaliana* Col-0 mitochondrial or plastid reference genome. Distribution of the genetic elements followed annotation information of the reference sequences. For detailed information about the positions of each cytosine and its annotation, see Supplementary Data Set 1. **D.** The number and the proportion of non-synonymous substitutions which occurs when tC is replaced with tT.

### Phenotypes and genotypes of T_1_ plants

We first examined the phenotypes of transformed T_1_ seedlings grown on sucrose-containing medium (Fig. 2A, B, Supplementary Table S1, Supplementary Fig. S3). Among the eight mitochondria-targeted (mt) WHY2-CD mutators, some or many of T_1_ individuals of all lines exhibited growth retardation phenotypes, except for the line G. mtCWC170_GSVG. In contrast, plastid-targeted (pt) mutators showed less pronounced growth retardation but displayed color-visible phenotypes such as whitening (A. ptCWC188), variegation (B. ptCWC170), and yellowing (E. ptGSVG). Several individuals with severe aberrant phenotypes, particularly those of A. mtCWC188, B. mtCWC170, E. mtGSVG, and A. ptCWC188, died before reproduction, producing seeds, resulting in no T_2_ progeny (Supplementary Table S1). For each mutator, a higher or comparable frequency of abnormal phenotypes was observed in mitochondria-targeted lines compared to plastid-targeted ones (Fig. 2B). These results suggest that WHY2-CD mutators targeted to mitochondria or plastids can introduce visibly distinct phenotypes, with varying degrees of severity.

**Fig. 2.**
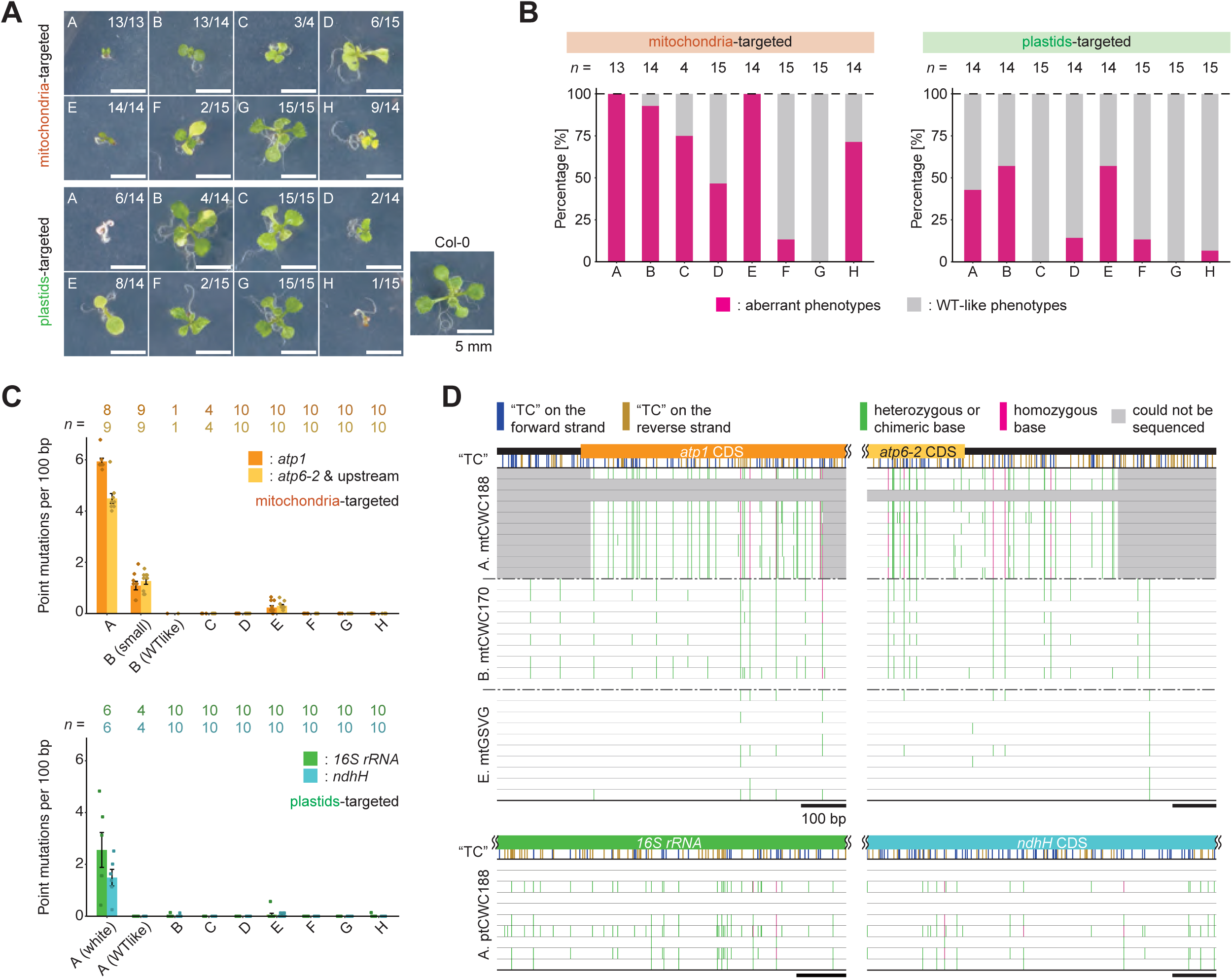
Phenotypes and genotypes of T_1_ plants. **A.** The severest phenotypes of T_1_ individuals among each construct. The fraction numbers show individuals of similar phenotypes/total germinated. 9 days after stratification (DAS). **B.** The ratio of T_1_ plants showing aberrant phenotypes. Phenotypic evaluation and photos are provided in Supplementary Table S1 and Supplementary Fig. S3. **C.** Point mutation rates in the regions Sanger-sequenced. Mean ± s.e.. **D.** Point mutation positions detected in individuals of mtCWC188, mtCWC170, mtGSVG, and ptCWC188.

To test whether WHY2-CD mutators induced mutations in organelle genomes, we PCR-amplified and Sanger-sequenced two short regions containing essential (*atp1* in mitochondria and *16S rRNA* in plastids) or non-essential (*atp6-2*^49^ in mitochondria and *ndhH*^50^ in plastids) genes. To avoid amplifying nucleus-localized highly homological mitochondrial DNA sequences (NUMTs^51, 52^), primers were designed to specifically target mitochondrial DNA (Supplementary Table S4). Even in these short regions (several hundred base pairs, Fig. 2C), C-to-T substitutions—defined as roughly ≥10% signal in Sanger chromatograms—were detected in A. mtCWC188, B. mtCWC170, E. mtGSVG, and A. ptCWC188. In B. mtCWC170 and A. ptCWC188, phenotypic data supported a link between mutations and abnormal growth. Mutation mapping (Fig. 2D) showed that (1) the number of mutations varied by construct, (2) not all “TC” sites were mutated, (3) some sites were mutated in multiple individuals and lines, and (4) some T_1_ plants had probable homoplasmic mutations. C-to-T changes in “tcC” contexts were also observed in A. mtCWC188. These results indicate that WHY2-CD mutators induced C-to-T mutations in the targeted organelle genomes with partial sequence preference.

### Next-generation sequencing (NGS) analysis of T_1_ plants

To assess the genome-wide mutagenic effects of WHY2-CD mutators in T_1_ plants, we performed short-read Illumina sequencing and single nucleotide polymorphism (SNP) call analysis. To determine whether C-to-T mutations occurred specifically in the targeted organelle genomes, bulk DNA samples from 5–11 T_1_ plants of mitochondria- or plastid-targeted lines D. E1347D and E. GSVG were sequenced, respectively. These four lines were selected for their distinct phenotypes (Fig. 2A, B; Supplementary Table S1) and sufficient biomass for sampling of DNA for NGS. In the E. mtGSVG sample, 6,441 C-to-T mutations above the threshold were detected in the mitochondrial genome, but two in the plastid genome (Fig. 3A; Supplementary Data Set 2). In the E. ptGSVG sample, 88 C-to-T mutations were identified in the plastid genome, and only two in the mitochondrial genome. In the D. mt/ptE1347D sample, no C-to-T mutations were detected in either of the organelle genomes. In all four samples, only a few C-to-T mutations were detected on nuclear chromosome 1, except near the centromeric region, where the abundance of repetitive sequences makes variant calling unreliable (Fig. 3A, Supplementary Fig. S4). These results indicate that C-to-T mutations were introduced almost exclusively in the organelle targeted by the corresponding signal peptide, at least when using the E. GSVG construct.

**Fig. 3.**
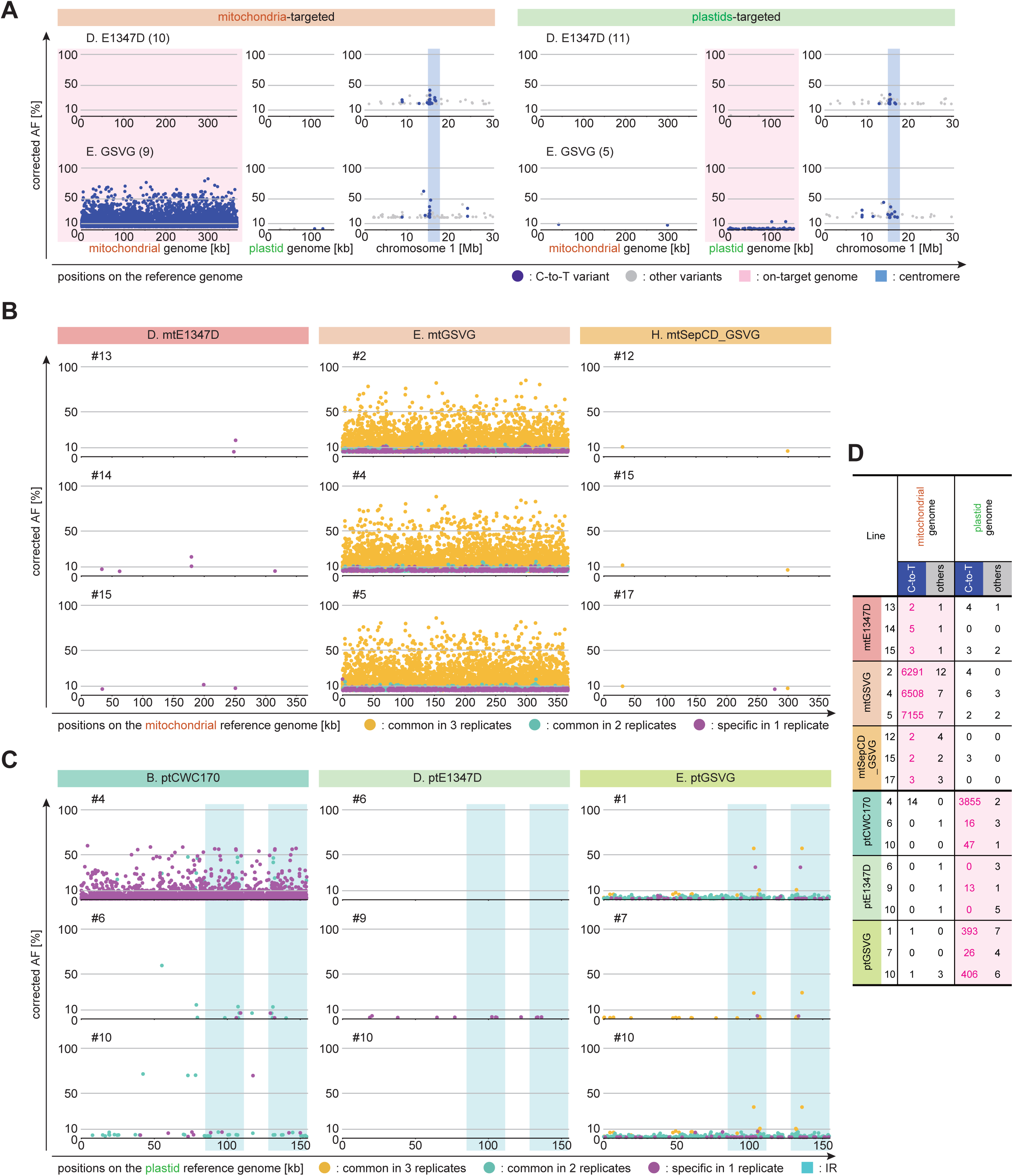
NGS and SNP call analyses of the whole organelle genomes of T_1_ plants. **A.** Corrected allele frequency (cAF) of each detected variant in bulk DNA samples. Numbers in parentheses represent the number of mixed individuals. The centromere region on chromosome 1 was defined by Naish *et al.* 2021^73^. **B, C.** cAFs of C-to-T mutations in three individuals per construct. Mutations common to three samples were colored yellow, those common to two samples were colored cyan, and those specific to one sample were colored magenta. **B.** T_1_ with mitochondria-targeted constructs, **C.** T_1_ with plastids-targeted constructs. The inverted repeat (IR) regions were defined by Sato *et al.* 1999^1^. **D.** The number of detected variants on the organelle genomes for samples **B** and **C.** On-target genome is colored magenta.

To examine the mutation profiles in individual T_1_ plants, three plants per line were independently sequenced (Fig. 3B–D). Samples were obtained from D. mt/ptE1347D and E. mt/ptGSVG (same constructs as in Fig. 3A), as well as from B. ptCWC170 and H. mtSepCD_GSVG, which showed visible abnormal phenotypes but retained fertility (Supplementary Table S1). In D. mtE1347D, two to five independent mutations were detected in the mitochondrial genome of each plant (Fig. 3B, D). The corrected allele frequencies (cAFs: sample AFs extracted by wild-type AFs at the same position) ranged from 5% to 15%, indicating heteroplasmy or chimerism involving mutated and wild-type mitochondrial DNAs. The mutation sites differed among the plants. In H. mtSepCD_GSVG, two to three mutations with cAFs between 5% and 10% were identified. Two of these mutations were common to all three plants (Fig. 3B, D). In E. mtGSVG, which was lethal or sterile, 6,200 to 7,200 mutations were detected across the mitochondrial genome (Fig. 3D). Most variants with cAFs greater than 15% were shared among the three plants (Fig. 3B). In addition, a structural change involving increased copy number in a specific mitochondrial region was observed in mtGSVG #2 (Supplementary Fig. S5A). This type of structural change in the mitochondrial genome resembles those reported in mitochondrial DNA repair mutants^53^. A strong preference for the “TC” sequence context, a known DddA target^47^, was confirmed (Supplementary Fig. S5B). A weak tendency for G or A at the –2 position was also observed, although this was not reported in Mok *et al.* 2020^47^. These results indicate that WHY2-CD mutators can introduce C-to-T mutations in the mitochondrial genome with partial sequence context specificity. In E. mtGSVG, excessive mutation levels may have contributed to lethality or sterility. In contrast, D. mtE1347D successfully generated viable T_1_ plants carrying distinct mitochondrial mutations.

For plastid-targeted lines, approximately 400 C-to-T mutations were detected in the plastid genomes of individuals #1 and #10 of E. ptGSVG, and 26 mutations in individual #7 (Fig. 3C, D). In D. ptE1347D, no mutations were found in two of the three individuals, while 13 mutations with cAFs around 1% were detected in individual #9. In B. ptCWC170, two individuals had approximately 50 mutations each, and individual #4 had 3,855 mutations (Fig. 3C, D). In terms of a high rate of cAFs, in E. ptGSVG, mutations with cAFs around 40% were observed only in individual #1. In B. ptCWC170, mutations with cAFs ≥ 50% were found only in individual #4. In these analyses, no homoplasmic mutations (cAF = 100%) were detected in any plastid- (and also mitochondria-) targeted samples (Fig. 3B, C). In B. ptCWC170 #4, the nucleotide preference surrounding mutated cytosines resembled that observed in E. mtGSVG, the preference for “TC” context (Supplementary Fig. S5B). These results indicate that C-to-T mutations were also induced in the plastid genome in T_1_ plants. The number and cAF distribution of these mutations varied among constructs and among individual plants. Both common and individually unique mutation sites were detected in plastid-targeted lines.

### Phenotyping of the T_2_ plants

At first, phenotypes of 40 T_2_ seedlings per T_1_ parental line were evaluated by visual observation. As a control, a recessive msh1 mutant line (CS3246; *msh1-2*) was used. To reset its mutated organelle genomes to intact at first, an F_1_ heterozygous msh1 mutant was generated by crossing *msh1-2* with the seed parent Col-0, five F_2_ plants homozygous for the *msh1* mutation were selected, and their F_3_ progeny were subjected to phenotypic analysis^40^. In these *msh1* F_3_ populations, approximately 25% of plants exhibited abnormal phenotypes such as variegation or growth retardation. In WHY2-CD mutator-transformed lines, abnormal phenotypes also appeared in T_2_ populations, but their frequency varied depending on the T_1_ parent, even within populations derived from the same construct (Supplementary Data Set 3, Supplementary Data Set 4, Supplementary Fig. S6). Approximately 0–50% of T_2_ plants from D. mtE1347D, H. mtSepCD_GSVG, B. ptCWC170, and E. ptGSVG showed abnormal phenotypes (Supplementary Data Set 3). Some individuals lacking seed GFP fluorescence, indicating the loss of the nuclear-integrated WHY2-CD construct through Mendelian segregation, also exhibited abnormal phenotypes (Supplementary Data Set 3, Supplementary Fig. S6). These findings suggest that WHY2-CD mutators can induce heritable organelle mutations with visible phenotypes, and that phenotypic screening in the T_2_ generation may be feasible for mutant screening, in a manner comparable to that observed in msh1 mutants-derived progenies.

### Genotyping of T_2_ plants

The sequenced T_1_ plants that produced T_2_ seeds did not exhibit homoplasmic mutations (Figs. 2, 3). However, previous studies have shown that heteroplasmic states in plant organelle genomes can become fixed to a single allele, homoplasmy, within one or two generations^17, 40, 54^. To evaluate this phenomenon in our system and assess the utility of each T_2_ population for the homoplasmic mutant screening, genotyping was implemented for T_2_ plants of B. ptCWC170 and E. ptGSVG, which had C-to-T mutations with cAFs ≥ 10% on the plastid genome in the T_1_ generation (Fig. 3C and Supplementary Table S1). Two plastid loci, G55202 and G102787 (Supplementary Data Set 2), which showed C-to-T variants with cAFs above 30% in T_1_ plants, were selected for analysis. Twenty T_2_ individuals with seed GFP fluorescence and 20 without GFP fluorescence were genotyped using PCR-Sanger sequencing. At the G55202 site, which results in a G82E amino acid substitution in the *rbcL* gene, six and five individuals homoplasmic for the mutated allele were identified in both GFP-positive and GFP-negative groups, respectively (Fig. 4A). While all individuals homoplasmic for the mutant allele (A) exhibited yellow-green leaves and growth retardation, 16 out of the total 18 individuals (three for GFP-positive and 15 for GFP-negative) homoplasmic for the wild-type allele (G) displayed wild-type-like phenotypes (Fig. 4B, C). Among GFP-positive seedlings, more than 10 showed heteroplasmic or chimeric genotypes. In contrast, no heteroplasmic or chimeric individuals were detected among GFP-negative seedlings (Fig. 4A). These results indicate that it is feasible to isolate homoplasmic plastid genome mutants in the T_2_ generation of B. ptCWC170 and E. ptGSVG, and that the mutation can be fixed in individuals in which the mutator construct was removed from the nucleus.

**Fig. 4.**
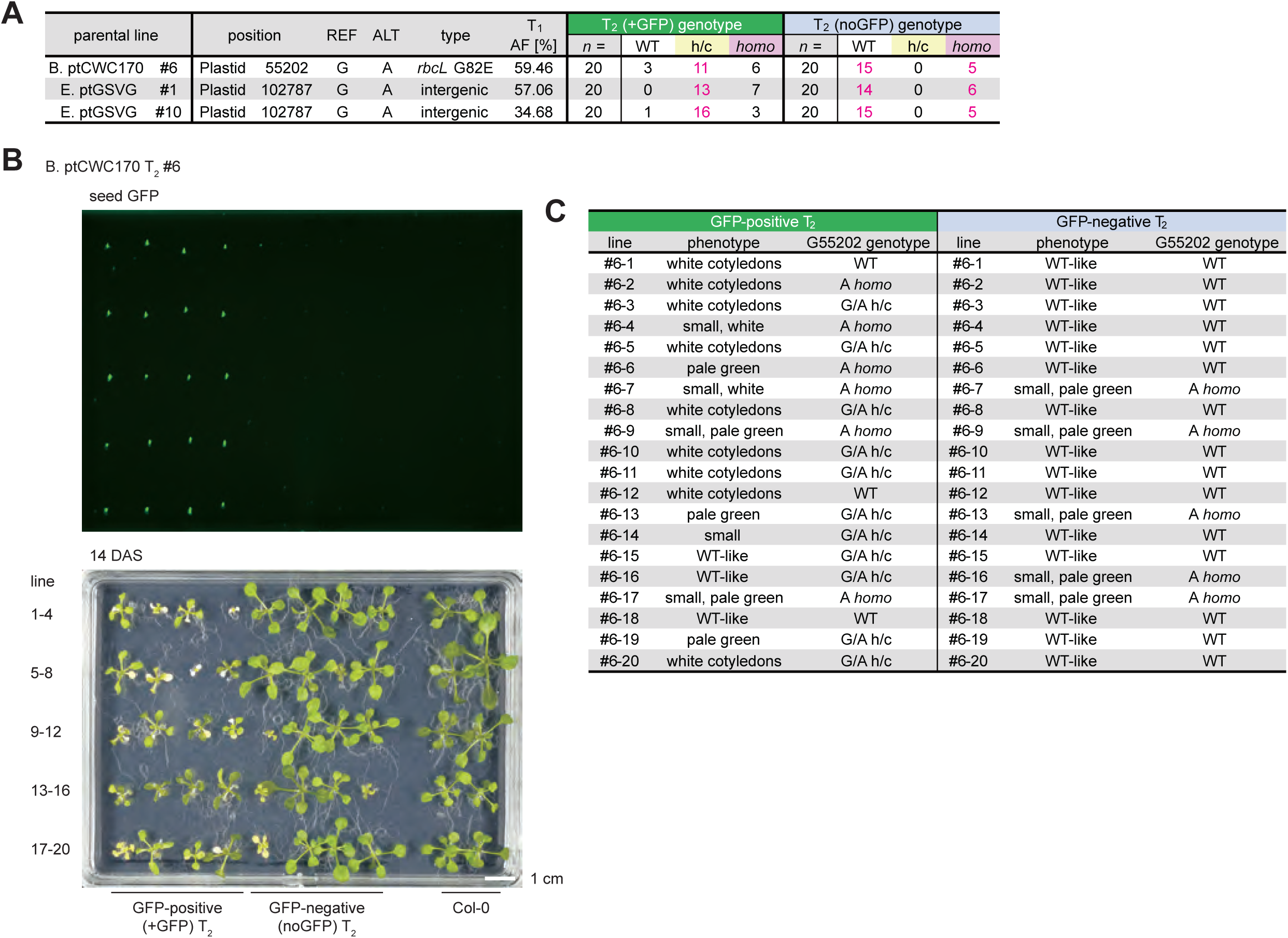
Genotypes of the plastid genome of B. ptCWC170, E. ptGSVG T_2_ plants. **A.** The genotypes of the same positions where C-to-T (G-to-A) variants were detected in NGS analysis of T_1_ (Fig. 3C, D), examined by Sanger sequencing. h/c: heterozygous or chimeric. **B.** Seed GFP fluorescence and phenotypes of the seedlings at 14 DAS. **C.** Phenotype description of each individual and its genotype of plastid genome G55202 analyzed by Sanger sequencing.

To investigate the number and positions of inherited C-to-T mutations across the plastid genome, whole-genome sequencing using Illumina short-reads was conducted on several GFP-negative T_2_ individuals derived from B. ptCWC170 #4 and E. ptGSVG #10. The T_1_ plant B. ptCWC170 #4 carried more than 1,000 heteroplasmic or chimeric mutations (Fig. 3D), whereas none of its three T_2_ progeny exhibited C-to-T mutations in the plastid genome (Supplementary Fig. S7). In contrast, all three T_2_ individuals from E. ptGSVG #10 carried mutations with corrected allele frequencies (cAFs) ≥ 99.5%, probably suggesting homoplasmic status (Fig. 5). The mutation sites differed among the three individuals, and the number of mutations appeared to vary by individual (Fig. 5B). The individual #10-24_noGFP, which showed an aberrant pale green to white phenotype, carried 13 homoplasmic C-to-T mutations, a higher number than those in #10-21_noGFP (one mutation) and #10-22_noGFP (one), which exhibited wild-type-like phenotypes (Fig. 5). Several C-to-T mutations were identified at positions not previously detected in the parental T_1_ plant (Fig. 5). These results indicate that 1) T_2_ individuals derived from E. ptGSVG can carry homoplasmic C-to-T mutations, and 2) they have independent, distinct mutations in individuals, and so suggest that 3) they are available for genome-wide mutant screening for the plastid genome mutants.

**Fig. 5.**
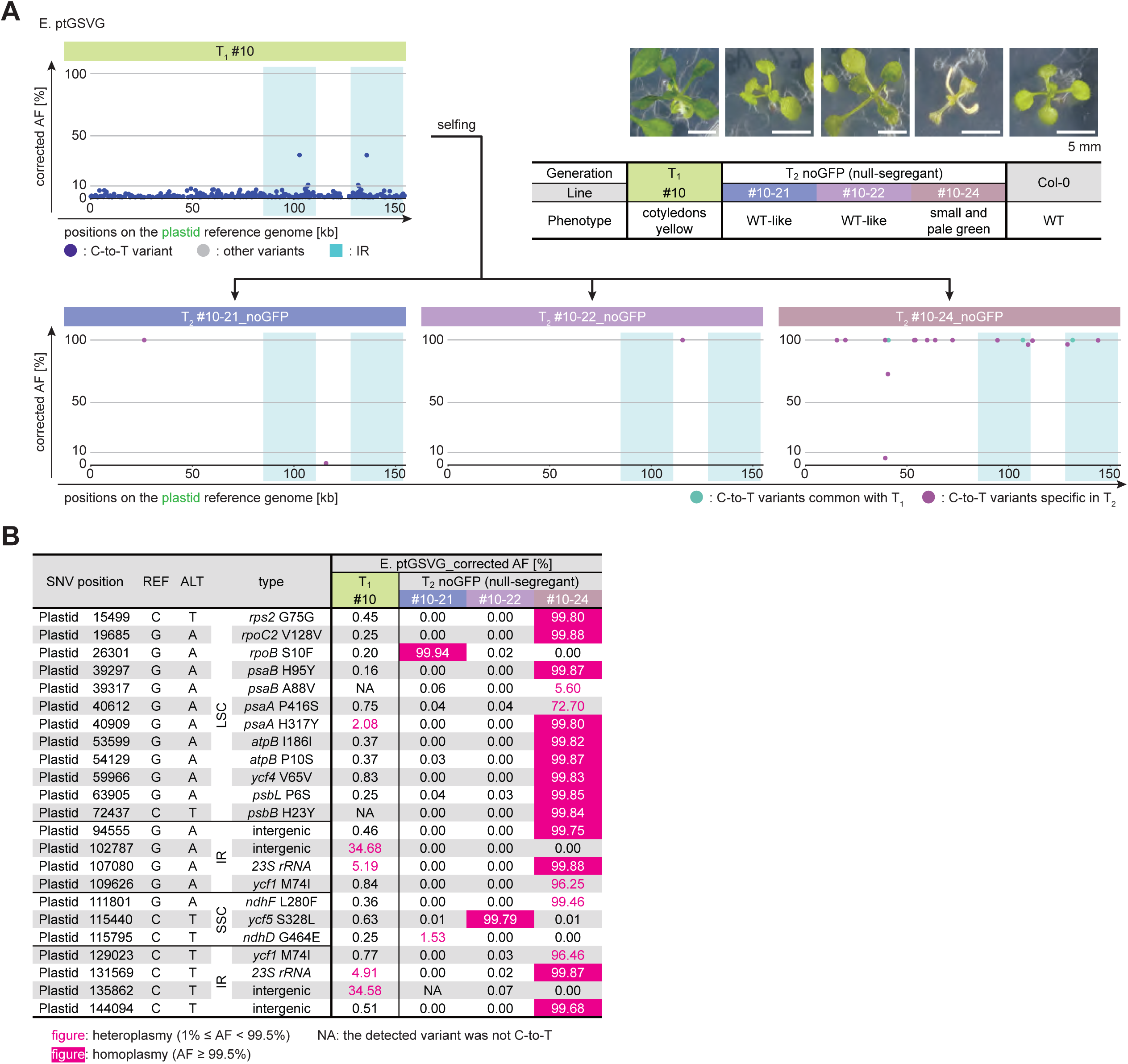
NGS and SNP call analyses of the whole plastid genome of E. ptGSVG T_2_ plants. **A.** Whole genome NGS and SNP call analysis of three T_2_ individuals derived from E. ptGSVG #10, and their phenotypes. Phenotypes are at 14 DAS. **B.** Positions of C-to-T variants with cAF and their genetic elements or consequent amino acid substitutions. NA means that the detected variant was not C-to-T. LSC, large single copy; IR, inverted repeat; SSC: small single copy.

### Identification of a mutation causing a chimeric, bleaching phenotype

Among T_1_ individuals transformed with B. ptCWC170, one individual exhibited a chimeric phenotype with distinct green and white tissues (Fig. 6A). To determine whether this phenotype was caused by a plastid genome mutation, DNA was extracted separately from each tissue and subjected to Illumina short-read sequencing. A specific C-to-T substitution (C78279T) with cAF of 98.47% was identified in the white tissue. This mutation introduced a premature stop codon in the plastid-encoded RNA polymerase (PEP) subunit A gene (*rpoA*), resulting in W204* (Fig. 6B). Both green and white tissues produced T_2_ seeds independently. Thirty T_2_ individuals derived from the green tissue and 15 from the white tissue were grown on 1/2 MS medium with sucrose (Fig. 6C). All progeny from the green tissue grew as well as the wild type, while all from the white tissue exhibited a fully white phenotype. Genotyping revealed that the green plants carried the wild-type W204 allele, whereas the white plants were homoplasmic for the W204* mutation (Fig. 6C). Although most white individuals failed to bolt or reproduce, one bolted but did not set seeds and ultimately died (Supplementary Fig. S8). These results suggest that the *rpoA* W204* mutation causes the bleaching phenotype but could be fertile and make seeds at least in the T_1_ generation, implying that PEP RNA polymerase is dispensable in bolting, flowering, reproduction, and embryogenesis, by supported by its chimeric green tissue. These also suggest the feasibility of isolating a lethal or severe phenotype-causing mutation from the chimeric parts.

**Fig. 6.**
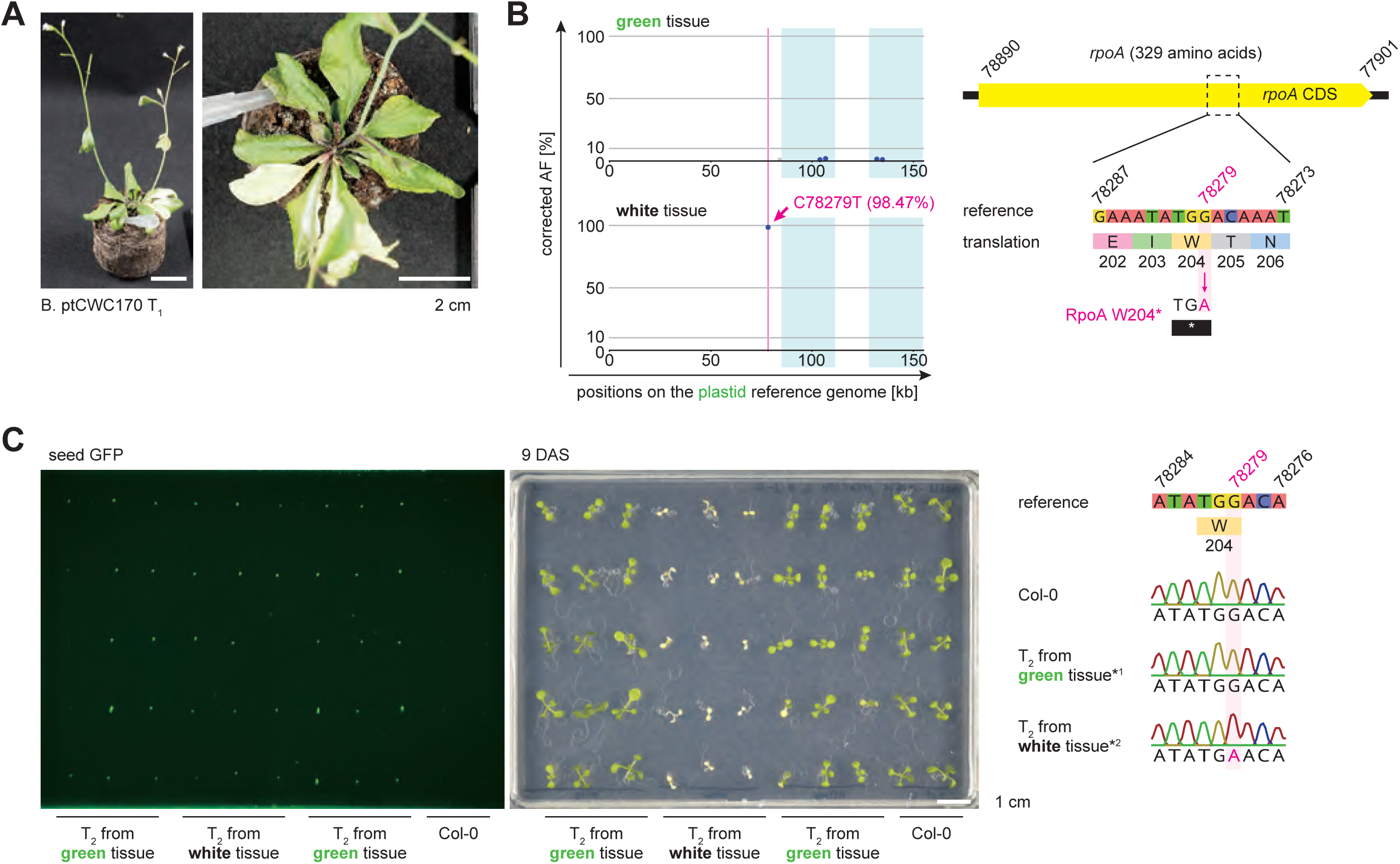
A mutation causing the white sector in a T_1_ plant and bleacing phenotype observed in the progeny. **A.** A phenotype of a chimeric T_1_ individual. 30 DAS. **B.** Scatterplot of SNP call results for the green and white tissues, respectively, and schematic diagram of the *rpoA* gene. **C.** Phenotypes of T_2_ individuals separately derived from green or white tissues. 9 DAS. Results of Sanger sequencing analysis of *rpoA* region. *1 and *2 are the representative results of eight individuals of green and white individuals, respectively.

### Selection of the mutant resistant to spectinomycin from the plastid-targeted T_3_ population

To assess the feasibility of large-scale screening for plastid genome mutants, spectinomycin (spm) selection was conducted on T_3_ populations of B. ptCWC170 and E. ptGSVG. Each T_3_ population was derived from one T_1_ line and subsequent 21 T_2_ plants grown in bulk. Several mutations in the plastid genome conferring spm resistance have been previously reported in five angiosperms, including two, C1015T and G1141A in 16S rRNA, in Arabidopsis (Supplementary Table S2; Refs. 13, 25, 33). Green plants emerged in two independent T_3_ populations derived from E. ptGSVG (#1 and #9) (Fig. 7A). Approximately 40 green individuals were observed in E. ptGSVG #1, and one in E. ptGSVG #9, out of approximately 7,500/plate spm-treated seedlings per population. To determine whether these resistant phenotypes were associated with base substitutions in the 16S rRNA gene, the corresponding region was amplified and sequenced in 12 individuals from E. ptGSVG #1 and one from E. ptGSVG #9. All resistant plants carried the same homoplasmic mutation, C102026T (C1015T in *16S rRNA* gene). This mutation was absent in three spm-sensitive T_3_ individuals (Fig. 7B). The positive control line had the same mutation (see Materials & Methods), but the possibility of seed contamination was excluded by detecting both the *16S rRNA* gene and the mutator transgene in a PCR assay (Supplementary Fig. S9). These findings demonstrate that plastid homoplasmic mutants conferring spm resistance can be isolated *de novo* from T_3_ populations generated by WHY2-CD mutators.

**Fig. 7.**
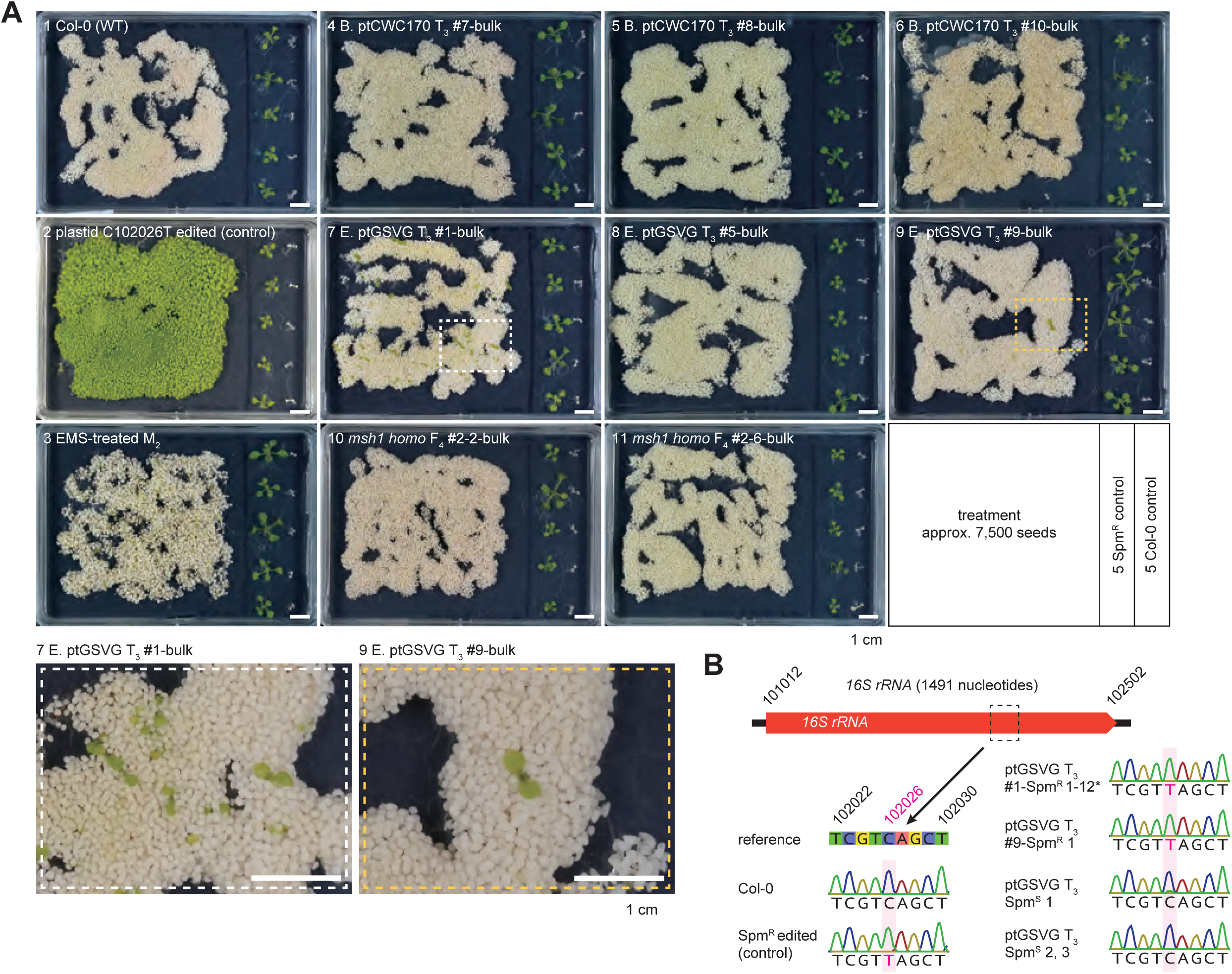
Spectinomycin screening treatment. **A.** Phenotypes at 14 DAS in the presence of spectinomycin (spm). Spm^R^, spectinomycin-resistant. **B.** Sanger sequencing analysis of *16S rRNA* gene. Asterisk: all twelve individuals examined had the genotype shown in the waveform. Spm^S^, spectinomycin-sensitive.

## Discussion

Organelle genomes have long been challenging to manipulate or improve through conventional breeding, largely due to their high copy numbers and non-Mendelian inheritance in most species. The advent of targeted base editing technologies enabling C-to-T and A-to-G substitutions^3, 11, 13–17, 55^ now allows precise and minimal modification of organelle genomes, opening opportunities for crop breeding outside GMO regulations. However, the loci that should be modified to achieve desirable traits remain largely unknown, with only a few examples—such as antibiotic or herbicide resistance and correlations between Rubisco large subunit variants and C4/C3 photosynthesis—currently documented^25–27, 56^. This knowledge gap contrasts with human mitochondrial DNA, where tens of thousands of natural single nucleotide variations and 117 pathogenic point mutations have been catalogued (www.mitomap.org, accessed Aug 2, 2025), enabling direct application of base editors for therapeutic and modeling purposes^47, 57^. Similarly, random mutagenesis tools and mitochondrial disease models are well established in animals, including mice^58, 59^ and Drosophila^60, 61^, leaving plant organelle research behind. Since crop improvements will be essential for yield enhancement, we aimed to develop plant organelle-genome-specific random mutagenesis as a forward genetic approach to efficiently improve the organelle genomes. In this study, we successfully achieved random mutagenesis biased toward C-to-T point mutations in Arabidopsis organelle genomes using the “WHY2-CD mutator.”

The C-to-T substitutions introduced by the WHY2-CD mutator appeared specific to either of the organelle genomes targeted by their N-terminal signal peptides (Fig. 3A), in contrast to chemicals or irradiation, which induce mutations across both nuclear and organelle genomes, and to msh1 mutants, which accumulate mutations in both plastids and mitochondria^40, 41^. Thus, the WHY2-CD mutator provides a tool for plastid- or mitochondrion-specific forward genetics. The introduced mutations, however, were not entirely random but showed a certain bias (Fig. 2D, Fig, 3B, C, Supplementary Fig. S5B). The preference for T at the −1 position of the substituted C is consistent with DddA characteristics^47^, yet our data suggest that not all “TC” contexts are equally prone to editing (Fig. 2D). Whether this reflects additional DddA sequence preferences or WHY2 binding properties remains unresolved. Nevertheless, useful C-to-T variants identified here can be validated through reverse genetics or transferred to related crops using ptp/mitoTALECD^13, 15^. Indeed, the isolated spm-resistant mutant carried an identical mutation at the position previously introduced by base editing (Fig. 7B), highlighting that although discovered in reverse order, such horizontal application is feasible.

To establish heritable and stable mutant lines, it is desirable that introduced mutations reach homoplasmy. We observed cases where heteroplasmic or chimeric mutations in T_1_ plants of B. ptCWC170 and E. ptGSVG fixed to homoplasmy in T_2_ (Fig. 4A). As *msh1* mutants show delayed segregation compared with wild type^54, 62^, mutations introduced by the WHY2-CD mutator in a wild-type background are expected to reach homoplasmy more rapidly than msh1-derived variants^40^. In three E. ptGSVG T_2_ null-segregants, the number of homoplasmic C-to-T mutations across per individual ranged from 1 to 13 (Fig. 5). With such numbers, promising variants could feasibly be validated reverse-genetically by generating single mutants via targeted base editing. Rapid segregation of heteroplasmic mutations within a generation agrees with previous reports^17, 54, 62^. However, it should be noted that the DNA extracted from limited tissue may either overrepresent homoplasmic sectors or miss variants entirely (Figs. 5, 6, Supplementary Fig. S7). Likewise, T_1_ genotypes represent only somatic mosaics and do not necessarily reflect the germline transmitted to all T_2_ progeny, as demonstrated by genotype differences among inflorescence branches leading to distinct T_2_ genotypes (Fig. 6). Despite these caveats, we show that, at least for the plastid genome, homoplasmic C-to-T mutants can be obtained within T_2_ or T_3_ generations, comparable to nuclear mutations (Figs. 4-7). For mitochondrial genomes, post-T_2_ segregation analysis was hindered by NUMTs, yet as suggested previously^54^, it is likely that homoplasmic mutants can also be recovered within 1–2 generations. Furthermore, the successful isolation of spm-resistant mutants through population screening (Fig. 7), identification of a chimeric *rpoA* knockout mutant among ∼30 ptCWC170 T_1_ plants (Fig. 6), and the correspondence between *rbcL* amino acid substitution (G82E) and pale-green, slow-growth phenotypes (Fig. 4B, C) collectively highlight the potential to isolate diverse mutants, including those affecting photosynthesis.

In summary, we demonstrate that (1) the WHY2-CD mutator introduces C-to-T mutations at multiple undefined sites specifically in the targeted organelle genome, (2) homoplasmic mutants can be isolated from T_2_ onward, and (3) diverse mutants with phenotypic alterations can be obtained through screening. Systematic screening of such populations will advance understanding of organelle gene and regulatory functions, while uncovering agriculturally valuable mutations in traits such as photosynthesis, herbicide resistance, and CMS. Combined with targeted base editing, this approach promises efficient direct engineering of organelle genomes, accelerating the realization of “organelle genome breeding.”

## Materials and Methods

### Plant Materials & Growth Conditions

*Arabidopsis thaliana* Columbia-0 (Col-0), *msh1* mutants [*msh1-2*; CS3372, Arabidopsis Biological Resource Center (ABRC)], and the other transgenic plants were grown at 22℃ under long-day conditions (16 h light, 8 h dark). Col-0 seeds were sown on 1/2 Murashige and Skoog (MS) medium (pH 5.7) containing 2.3 g/L of MS Plant Salt Mixture (Wako), 500 mg/L of MES, 10 g/L of sucrose, 1 mL/L of Gamborg’s Vitamin Solution (Sigma–Aldrich) and 6 g/L of agar. Medium was supplemented with 125 mg/L Claforan for the transgenic T_1_ plants. Most seedlings, at 2–3 weeks old, were transplanted to Jiffy-7 (Jiffy Products International) and subsequently subjected to Agrobacterium transfection.

### Vector Construction

The nucleotide sequences encoding CD & UGI coding regions were the same as the ones used in ptpTALECD^13^. The coding regions of WHY2-CD (wild type)-UGI and GSVG variant of CD^48^ were artificially synthesized (EurofinsGenomics) with codon optimization for A. thaliana. The E1347D variant of CD is an attenuated strain isolated during cloning of wild-type CD. We tried to clone Ti plasmids containing full-length DddA, but it was unsuccessful (Supplementary Fig. S1). For A. CWC188, B. CWC170, F. CWC188_GSVG, G. CWC170_GSVG, in order to avoid the cleavage of the WHY2-native organelle targeting signal, possibly resulting in separation and consequent inactivation of the mutator, the N-terminal 50 or 68 amino acids of WHY2 were removed based on cleavage prediction by TargetP 2.0 (https://services.healthtech.dtu.dk/services/TargetP-2.0/) (Supplementary Fig. S10). The entry vectors for C. SepCD were cloned by In-Fusion reaction using ptpTALECD entry vectors. The entry vectors for D. E1347D, E. GSVG were cloned by In-Fusion reaction using synthetic WHY2-CD (wild type)-UGI. The entry vectors for H. SepCD_GSVG were cloned by In-Fusion reaction using C. SepCD entry vectors. The reading frames in the assembled first and third entry vectors and the second entry vector (below) were transferred into the Ti plasmid by multi-LR reaction with LR Clonase II Plus (Thermo Fisher Scientific). The backbone of destination vector was pK7WG2^74^. RPS5Apro cloned from pKAMA-ITACHI1_1R^75^ was integrated as promoter^49, 76^. Either N-terminal 36 amino acids of ATPase δ prime subunit^46^ as MTP or N-terminal 51 amino acids of RecA1 (At1g79050)^13^ as PTP were also integrated. *Ole1* pro:: Ole1-GFP derived from pFAST02^77^ was inserted for selection of transformed seeds. The second entry vector harboring Arabidopsis heat-shock protein terminator, *RPS5A*pro and MTP/PTP was used for Ti plasmid construction of C. SepCD and H. SepCD_GSVG. Single-LR reaction was performed to construct D. E1347D and E. GSVG. Ti-plasmids of A. CWC188, B. CWC170, F. CWC188_GSVG, G. CWC170_GSVG were directly cloned by In-Fusion.

### Plant Transformation

Plant transformation methods and selection of transgenic T_1_ seeds were described previously^13^. Each of the vectors was introduced into *A. tumefaciens* strain C58C1, then transformed into Col-0 by floral dipping^63^. Transgenic T_1_ seeds were selected by observing seed GFP fluorescence.

### Plant Phenotyping

Arabidopsis phenotypes were observed at 9 or 14 DAS. The phenotypes were evaluated by their appearance, then recorded by digital cameras, EM-5 (Olympus) or α6400 (SONY).

### Plant Genotyping

DNAs were roughly extracted from an emerging true leaf or cotyledon of the seedlings at 14-21 DAS, boiled in DNA extraction buffer (100 mM Tris HCl & 5 mM EDTA) at 98℃ for 15 min. PCRs were performed using the primers listed in Supplementary Table S4 with KOD One PCR Master Mix or Quick Taq HS DyeMix (TOYOBO). Sanger sequencing was performed by EurofinsGenomics or FASMAC, and the results were analyzed by Geneious Prime 2025.02. (Biomatters).

### NGS Analysis

Total DNAs were extracted from approximately 3-4-week-old, 3-4 Arabidopsis rosette leaves (except for the samples marked with * in Supplementary Data Set 2), by DNeasy Plant Pro Kit (QIAGEN) or Isospin Plant DNA (NIPPON GENE) with standard protocols. For analysis shown in Fig. 3A, paired-end libraries using TruSeq DNA PCR-Free (350) (Illumina) and sequencing of 4 Gb/sample using NovaSeq6000 (Illumina) platform were carried out by PAGS (https://www.genome-sci.jp). For analyses shown in Fig. 3B and below, paired-end libraries using Nextera DNA Flex Library Prep Kit (Illumina) and sequencing of 4 Gb/sample using NovaSeq6000 or NovaSeqX (Illumina) platform were carried out by MacrogenJapan.

Paired-end 150-bp reads were trimmed by fastp v0.23.4^64^ and then mapped to the reference genomes by BWA v2.2.1^65^. We filtered out inadequate mapped reads with mapping identities ≤ 97% or alignment cover rates ≤ 80% or supplementary flags by samtools v1.20^15, 66^. SNPs were then called by samtools mpileup (-uf -d 50000 -L 2000 for organelle genomes, -uf -d 1000 -L 2000 for chromosome 1) and bcftools (v1.20) call (-m -A -P 0.1.) SNPs with allele frequency (AF) of; 1. AF in the same position of the wild-type sample was below the threshold (mitochondria: 0.5%, plastids: 0.5%, chromosome 1: 10%), and 2. sample AF correction by the wild-type sample (cAF) was over the threshold (mitochondria: 5%, plastids: 1%, chromosome 1: 20%) were finally counted. Furthermore, SNPs on chromosome 1 were counted only when their read depths were ≥ 15 and ≤ 50 (Supplementary Fig. S11). The reference genomes used: BK010421.1 for mitochondrial genome^67^, AP000423.1 for plastid genome^1^, CP002684.1 for chromosome 1^68^, CP002685.1 for chromosome 2^69^, CP002686.1 for chromosome 3^70^, CP002687.1 for chromosome 4^71^, and CP002688.1 for chromosome 5^72^.

### Spectinomycin Screening

We used six independent T_3_ populations, each established from 21 randomly selected T_2_ plants derived from one T_1_ individual. To prepare one population, 21 T_2_ plants were grown and their seeds were harvested in bulk (Supplementary Fig. S12). Approximately 250 µL of seeds per T_3_ population (about 7,500 grains) were sown on 1/2 MS medium supplemented with 20 mg/L spm. For control, a null-segregant T_4_ mutant population generated by targeted plastid genome editing, then harboring the homoplasmic mutation conferring spm resistance (plastid C102026T^13^), and Col-0 were used. Three msh1 mutant F_4_ populations and an EMS mutant M_2_ population were also tested.

### Image Processing

Plant images were taken by digital cameras, EM-5 (Olympus) or α6400 (SONY). Gel images were taken by ChemiDoc MP Imaging System (BioRad). They were processed with Adobe Photoshop 2025 (Adobe). Figures and tables were made with Adobe Photoshop 2025, Adobe Illustrator 2025 (Adobe), Microsoft Excel (Microsoft), R 4.4.0, and Python 3.11.

### Accession Numbers

Sequence data from this article can be found in the GenBank/EMBL data libraries under accession numbers: BK010421.1 (mitochondrial reference), AP000423.1 (plastid reference), CP002684.1 (chromosome 1 reference), CP002685.1 (chromosome 2 reference), CP002686.1 (chromosome 3 reference), CP002687.1 (chromosome 4 reference), CP002688.1 (chromosome 5 reference), At1G71260 (WHY2), AtMG01190 (atp1), AtMG01170 (atp6-2), AtCG00920 (16S rRNA), AtCG01110 (ndhH), At3G24320 (MSH1), AtCG00490 (rbcL), and AtCG00740 (rpoA).

## Acknowledgments

We thank Ms. Yoshiko Tamura, Ms. Reiko Masuda, and Dr. Hideki Takanashi for their technical support. The *msh1-2* seeds were kindly provided by the Arabidopsis Biological Resource Center.

## Author Contributions

Y.H., I.N., N.K. and S.A. initiated and designed the project. N.K. and Y.H. constructed the vectors and performed the main part of this research. N.K., M.O., T.I. and S.A. analysed next-generation sequencing data. N.K. and S.A. wrote the manuscript.

## Supplementary Information

**Supplementary Fig. S1.**
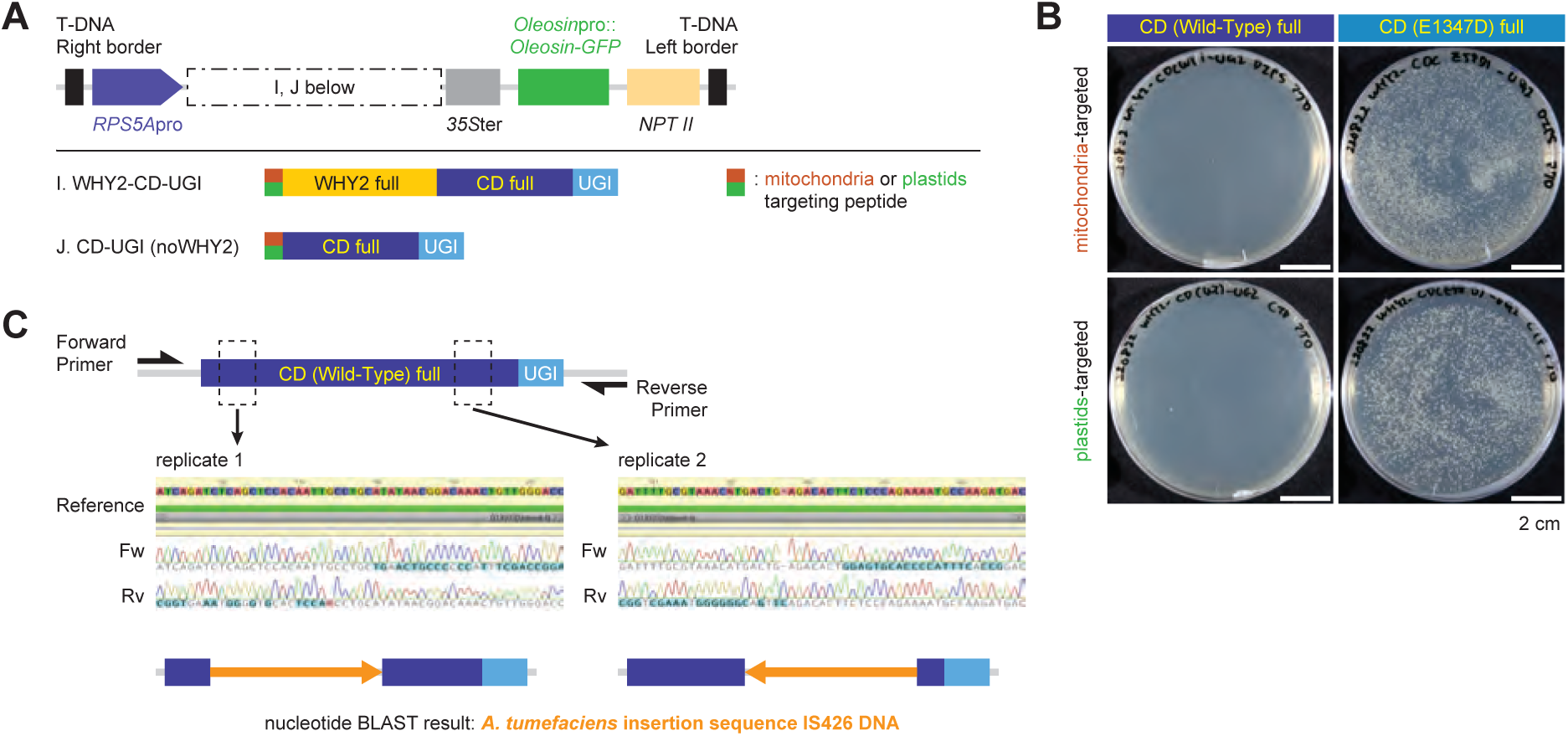
Obstacles encountered during the design and construction of the WHY2-CD mutator (supporting Fig. 1) **A.** A schematic illustration of WHY2-CD mutator containing full-length, wild-type DddA as cytidine deaminase (CD). **B.** Transformation results of *Agrobacterium* with constructs expressing wild-type, full-length CD. Efficient transformation was not achieved. As a control, a weakly toxic/low-activity CD variant (E1347D) was transformed at the same DNA concentration. As a result, *Agrobacterium* transformation of only J. ptCD-UGI was successful and one of its clones was subsequently subjected for Arabidopsis transformation. **C.** Sanger sequencing results of J. ptCD-UGI region in two Arabidopsis T_1_ plants. Transposon insertions were detected in the CD domain, resulting in inactivation of CD and base substitution activity. These results indicate that cloning and introducing a functional, wild-type full-length CD into *Arabidopsis thaliana* is technically difficult.

**Supplementary Fig. S2.**
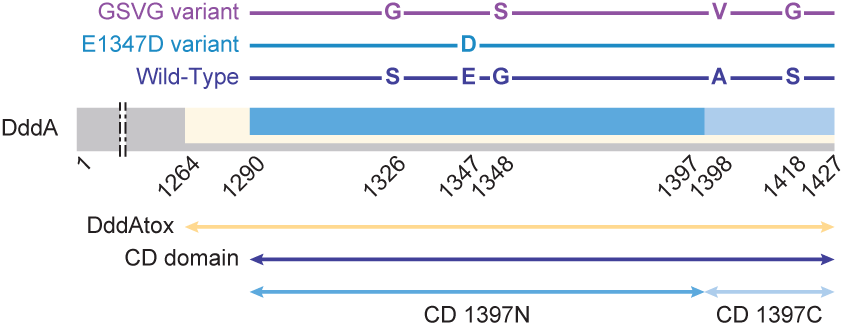
Amino acid structure and variants of the cytidine deaminase, Double-stranded DNA deaminase toxin A (DddA) used in this study (supporting Fig. 1)

**Supplementary Fig. S3.**
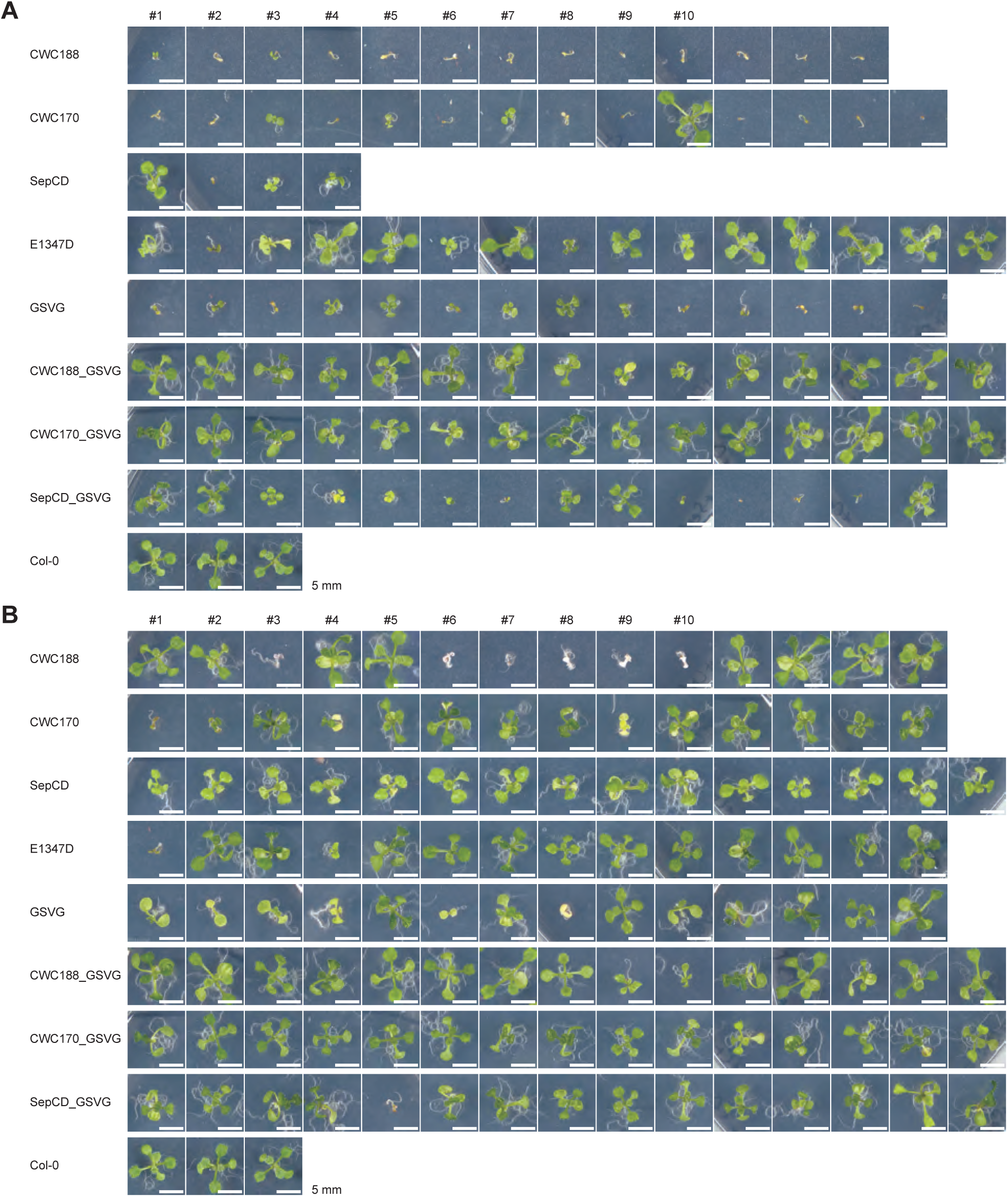
Photographs of all T_1_ individuals used for phenotyping in Fig. 2B (supporting Fig. 2) Individuals labeled with #1-#10 were subjected to Sanger sequencing analysis in Fig. 2C and 2D. **A.** Plants transformed with the mitochondria-targeted constructs. **B.** Plants transformed with the plastid-targeted constructs.

**Supplementary Fig. S4.**
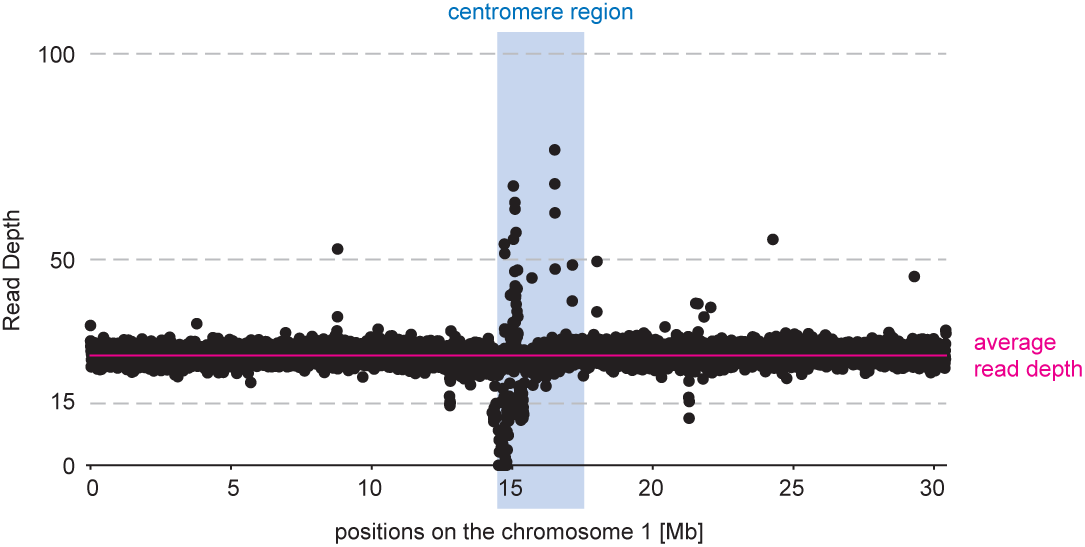
Read depths on the chromosome 1 of the sequencing data of the Col-0_bulk sample (supporting Fig. 3) This graph shows the abundance of repetitive sequences in the centromeric region, which makes variant calling difficult. For more detailed sample information, see Supplementary Data Set 2.

**Supplementary Fig. S5.**
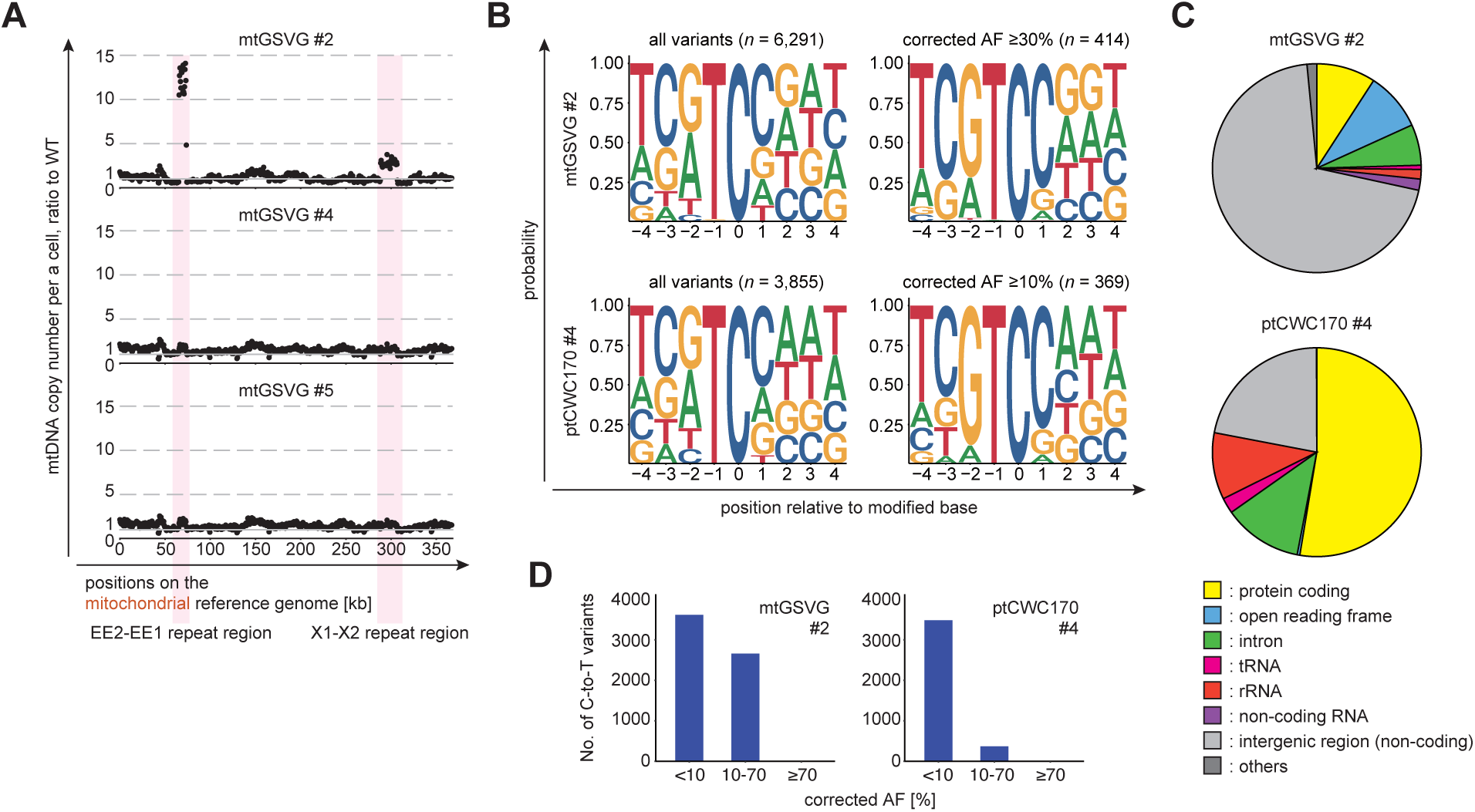
Additional analyses on NGS data of T_1_ individual samples (supporting Fig. 3) **A.** Depth of mitochondria-mapped reads of the three samples from E. mtGSVG, plotted by 500 bp. Vertical axis represents ratio of read depth to wild-type sample. Panels **B-D** are analyses with mtGSVG #2 and ptCWC170 #4 sequencing data, where more than 3,000 C-to-T mutations were detected on the on-target organelle genomes (Fig. 3D). **B.** Probability sequence logo of the region flanking mutated cytosines. ±4 positions of variant-detected cytosine are shown in the panel. No noteworthy base frequency bias was observed farther than ±4. **C.** The proportion associated with each genetic element of the variant-detected cytosines on the on-target genome. **D.** The number of C-to-T variants detected, categorized by corrected allele frequency (cAF) as less than 10%, between 10% and 70%, and greater than 70%.

**Supplementary Fig. S6.**
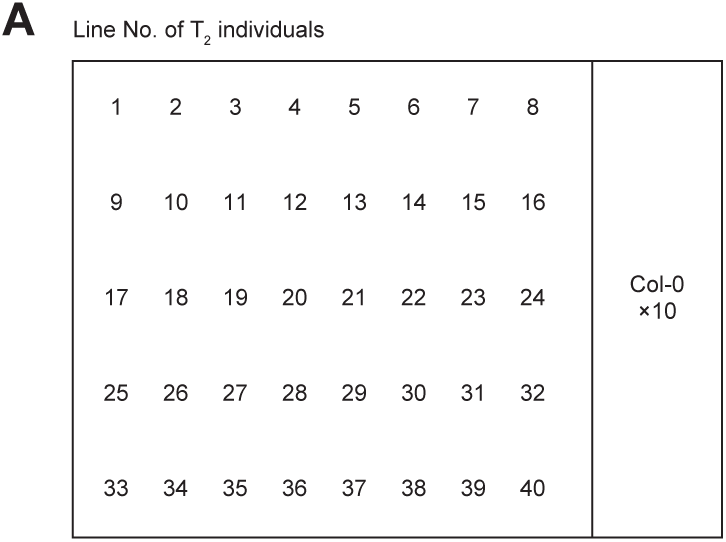
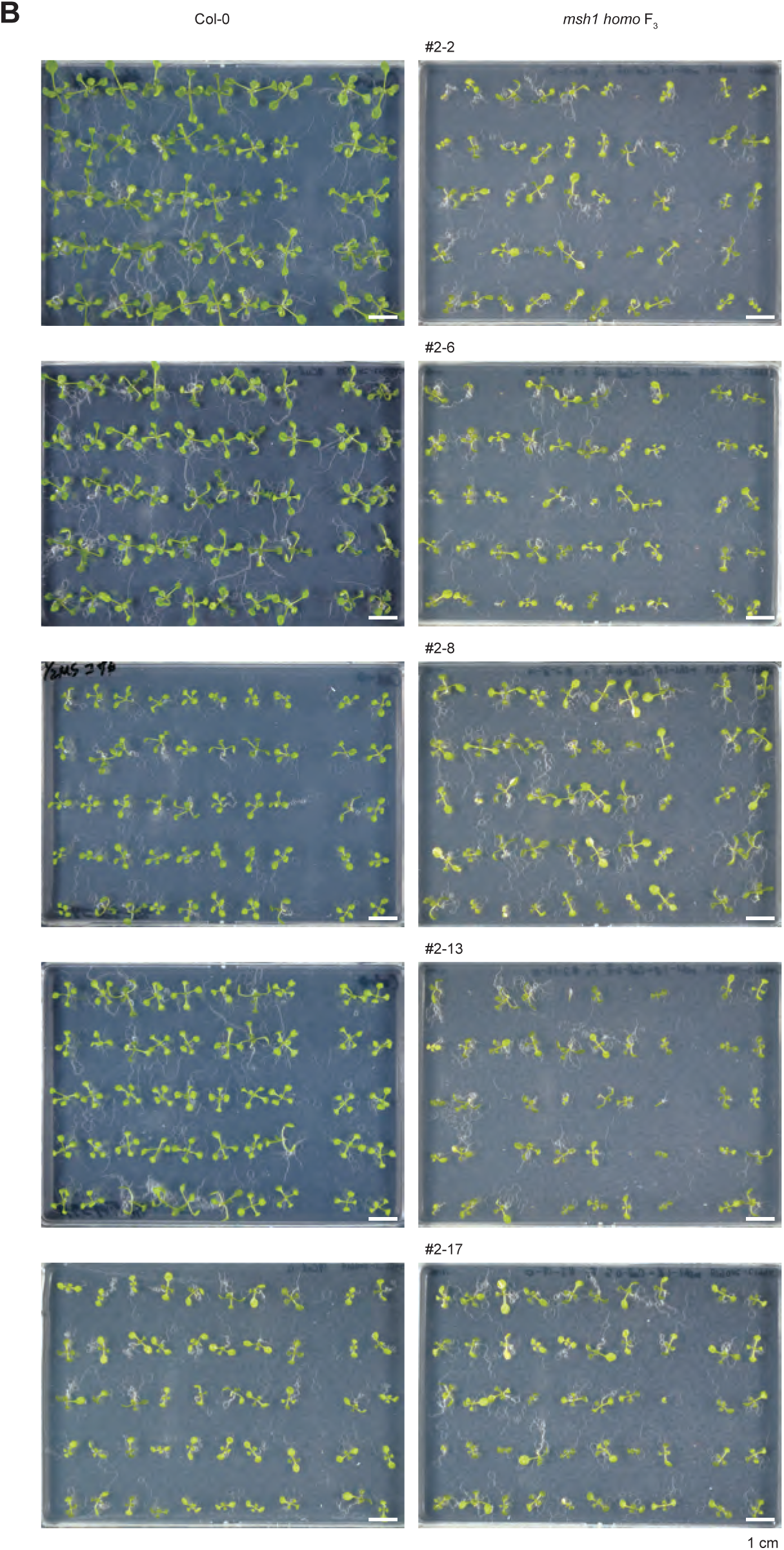
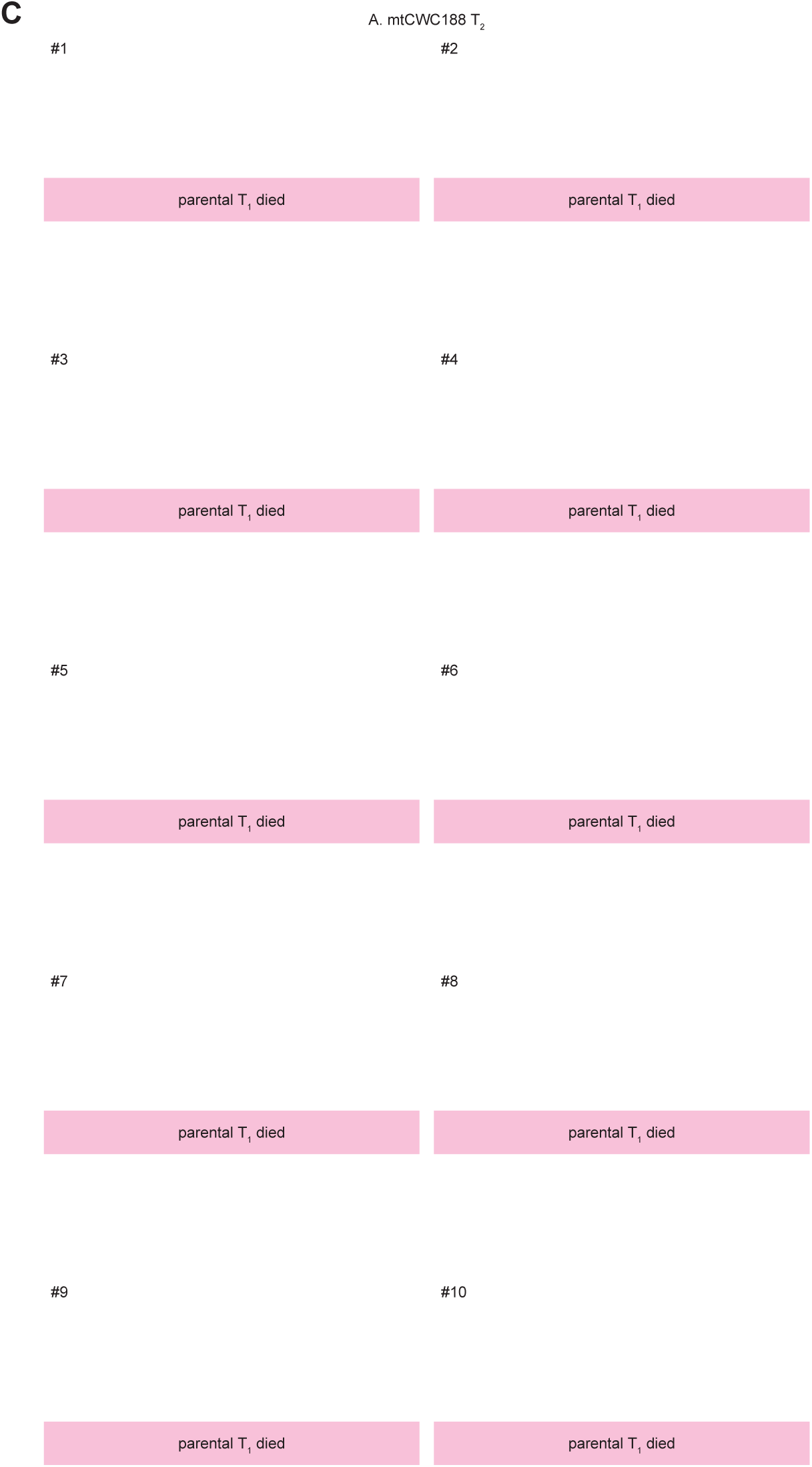
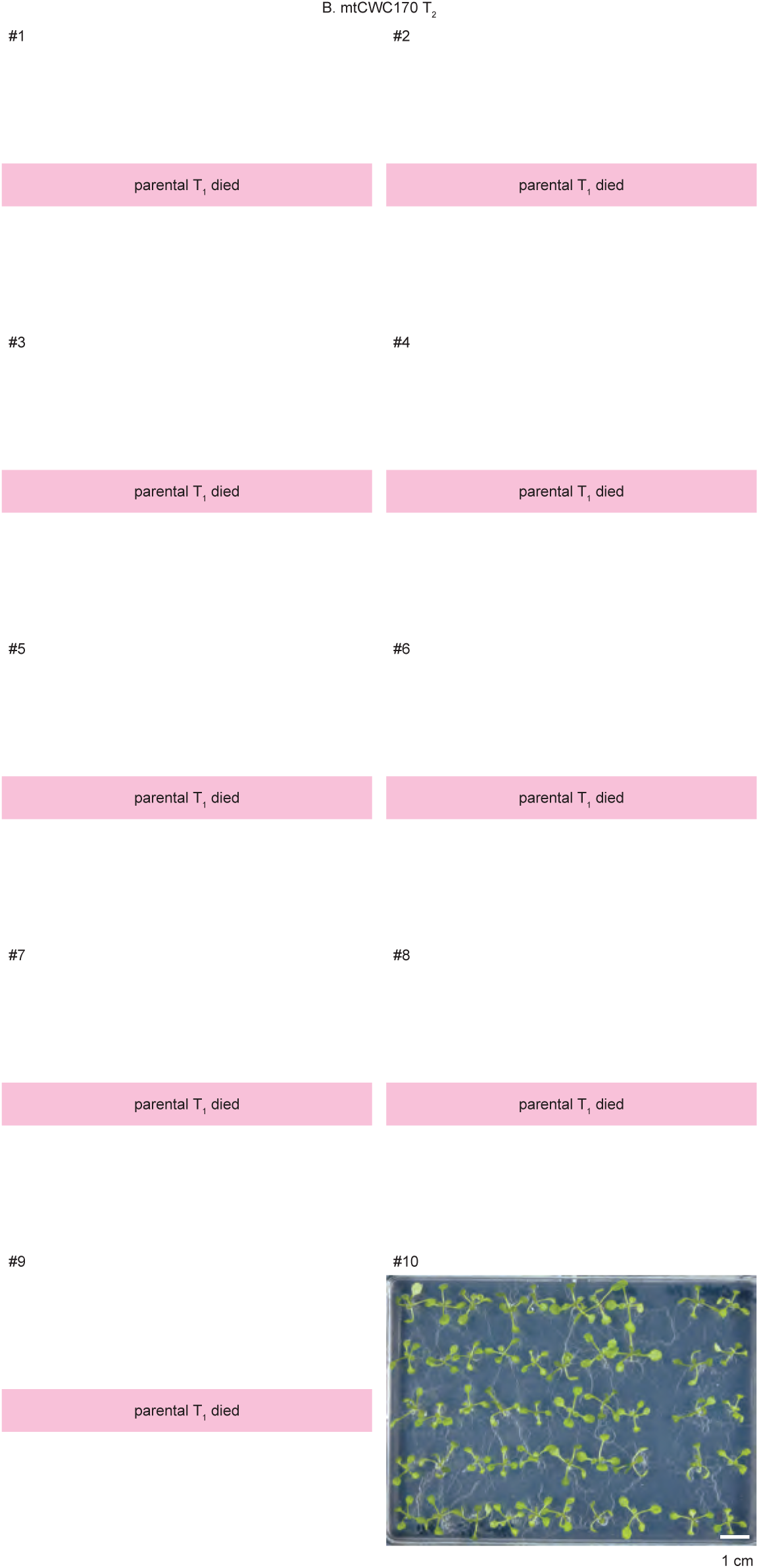
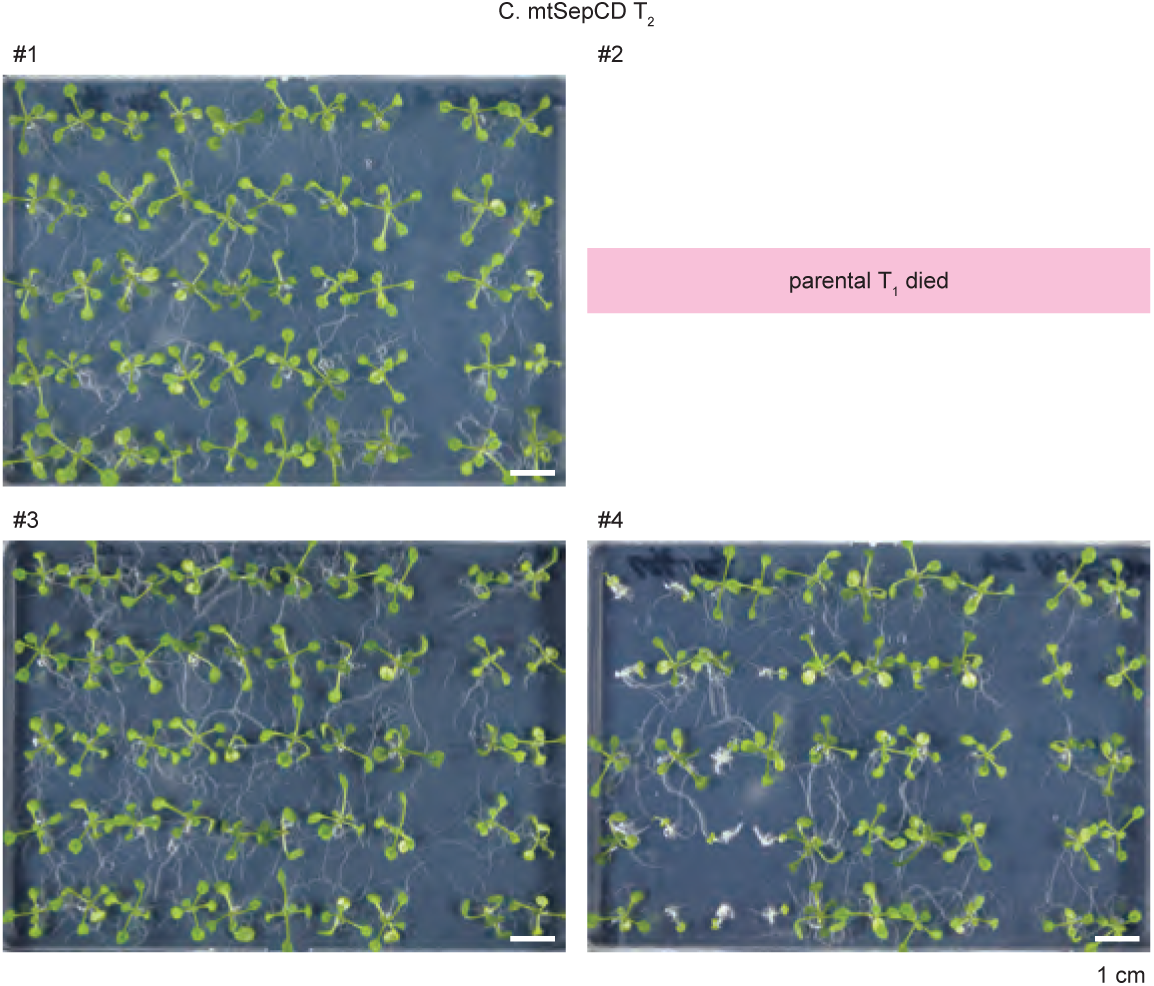
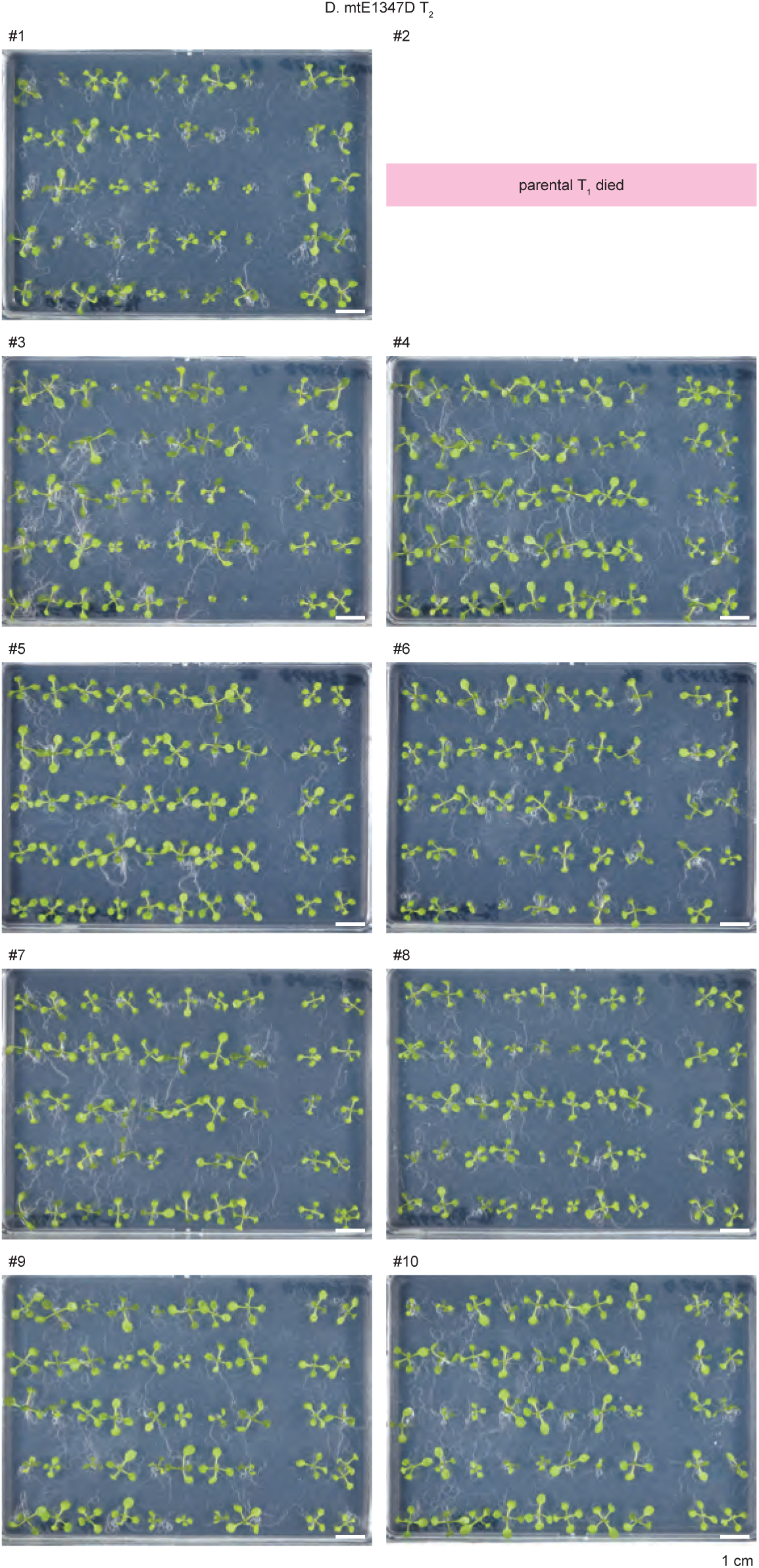
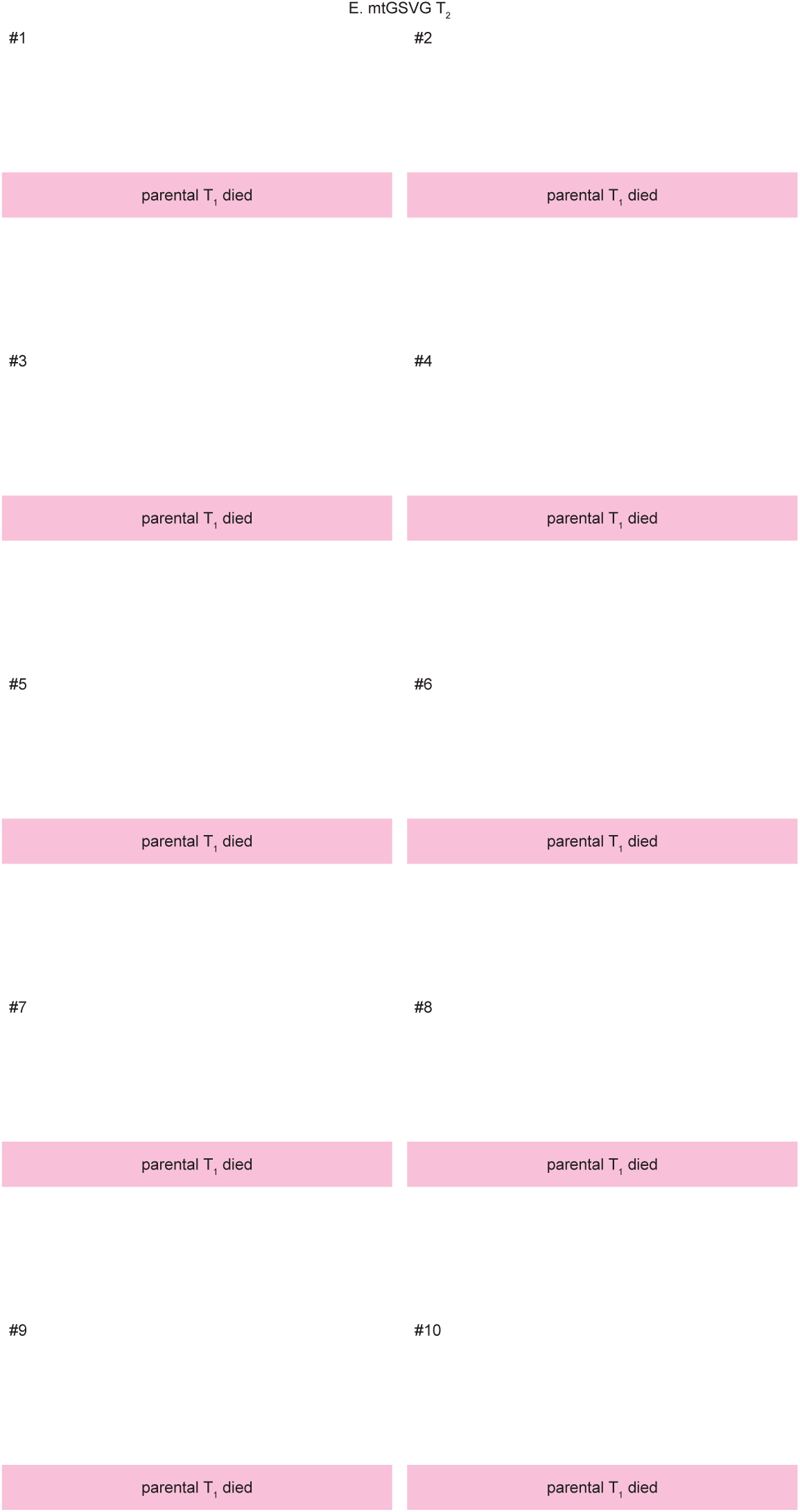
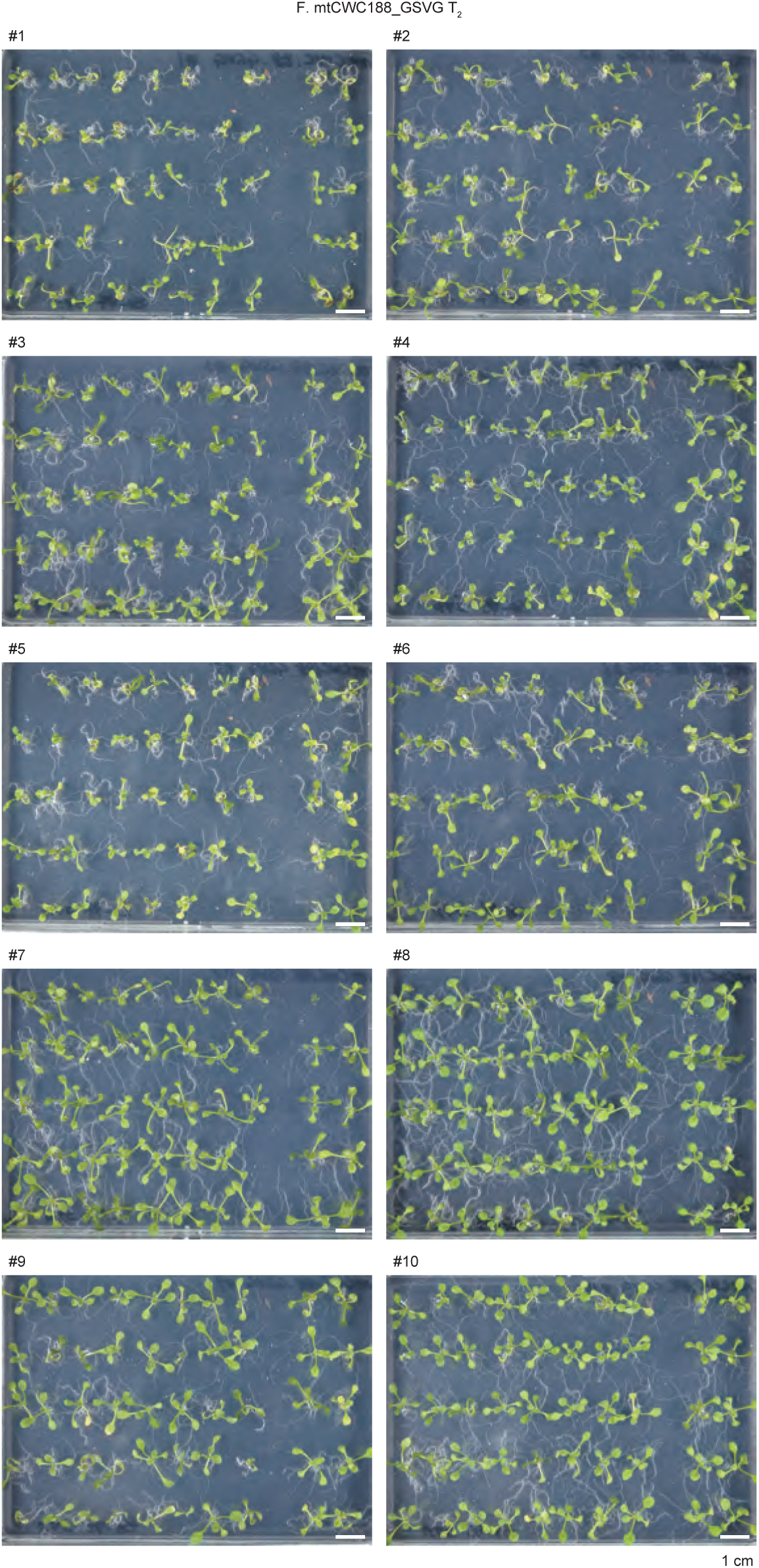
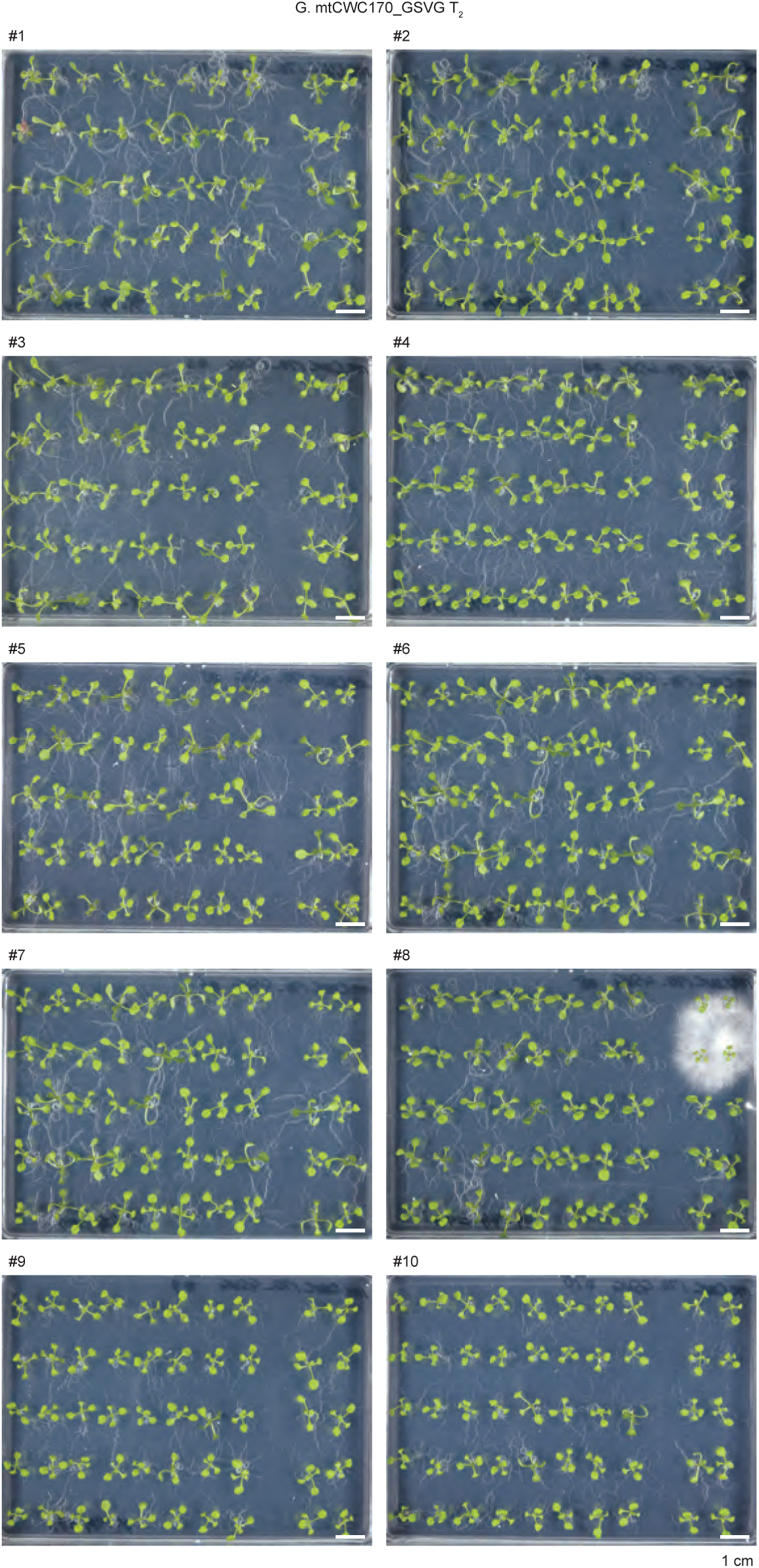
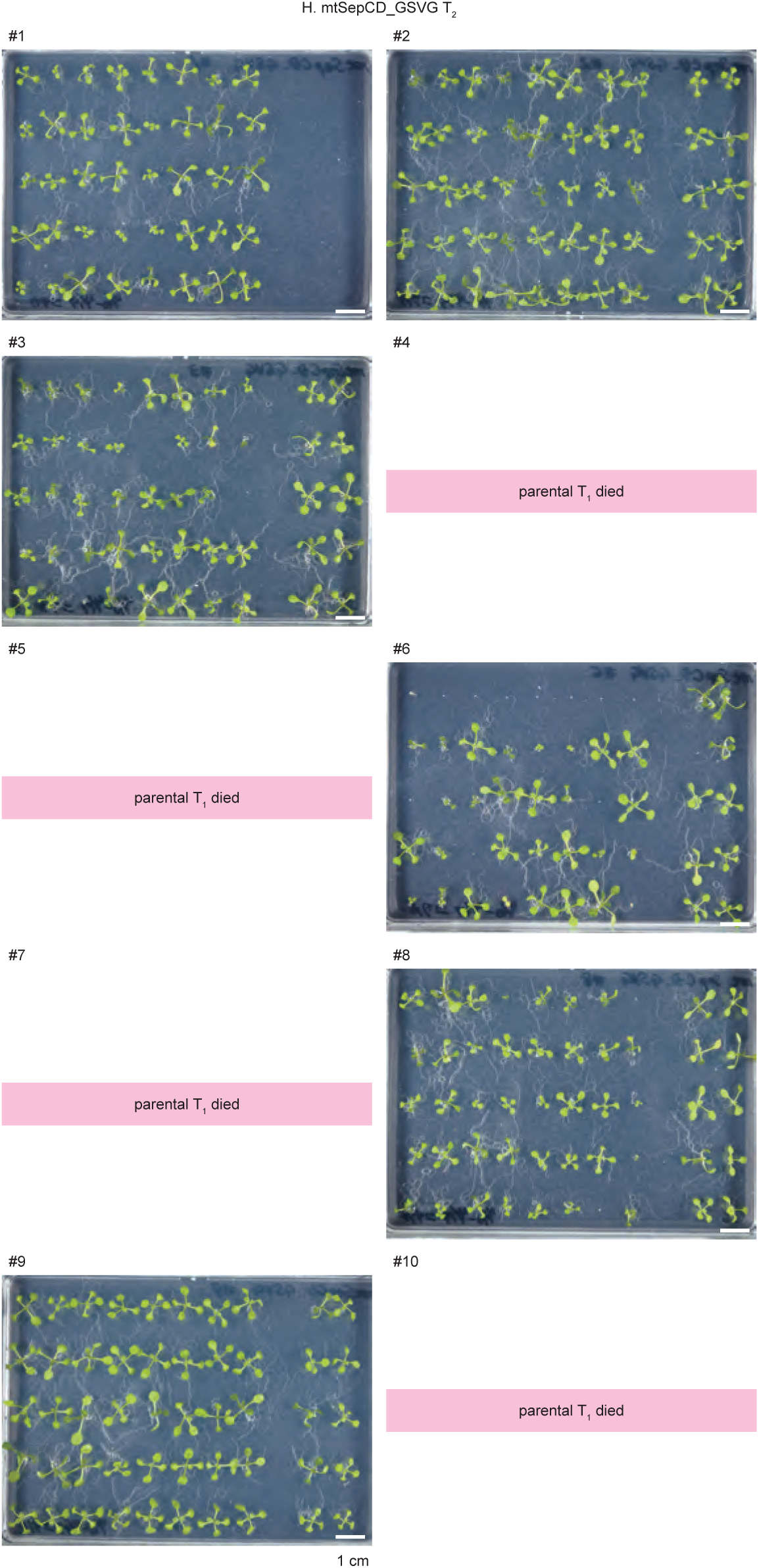
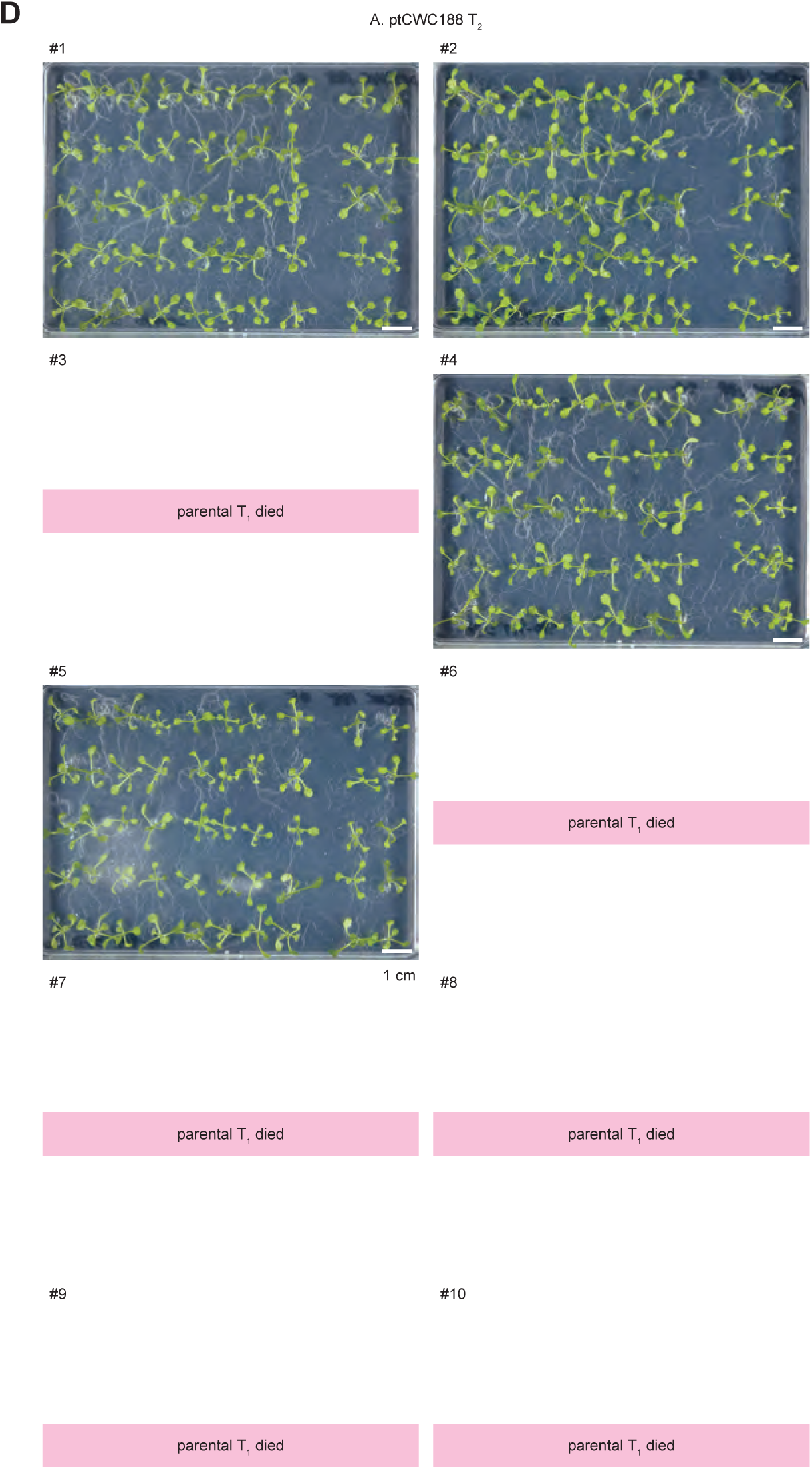
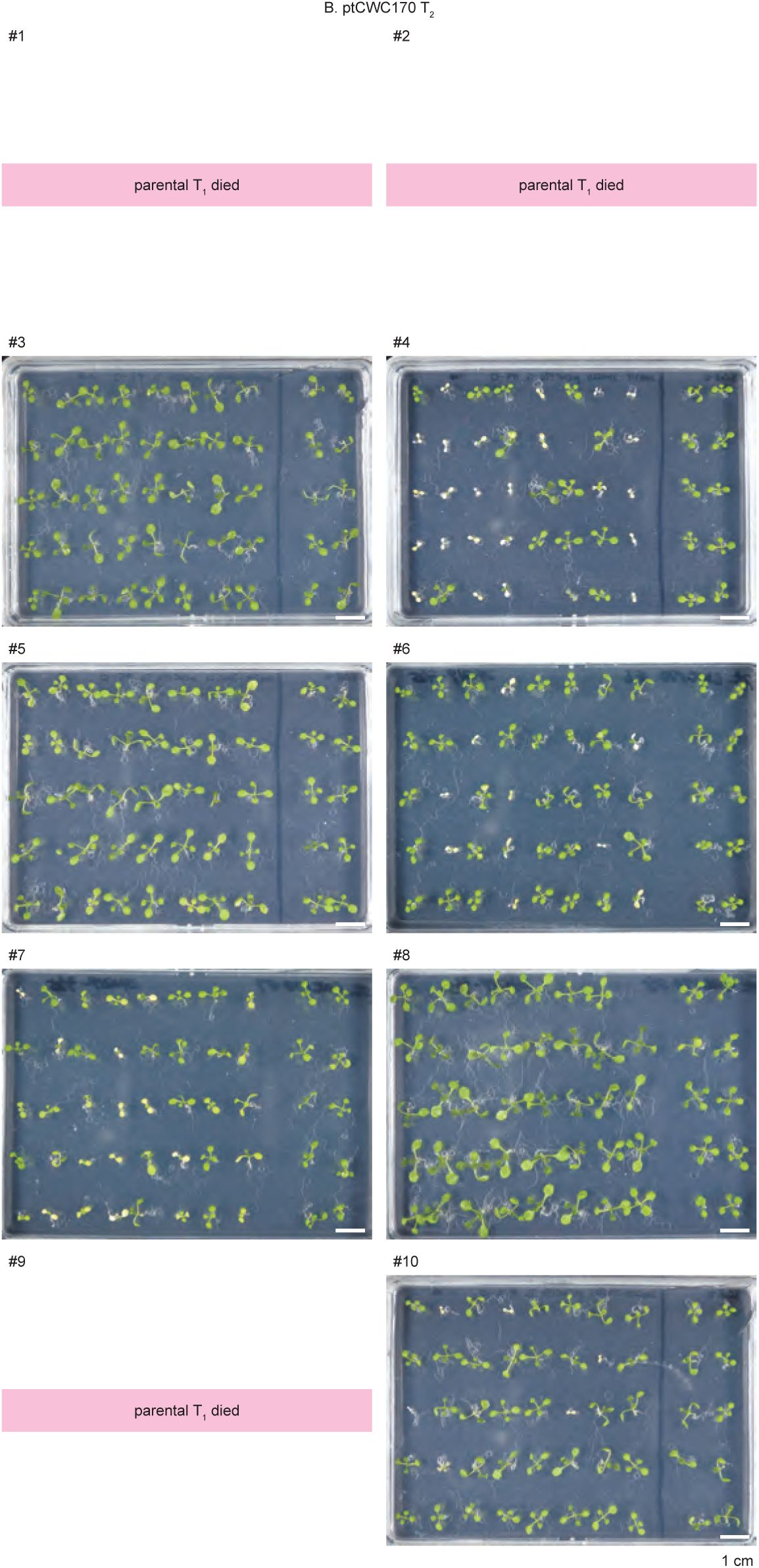
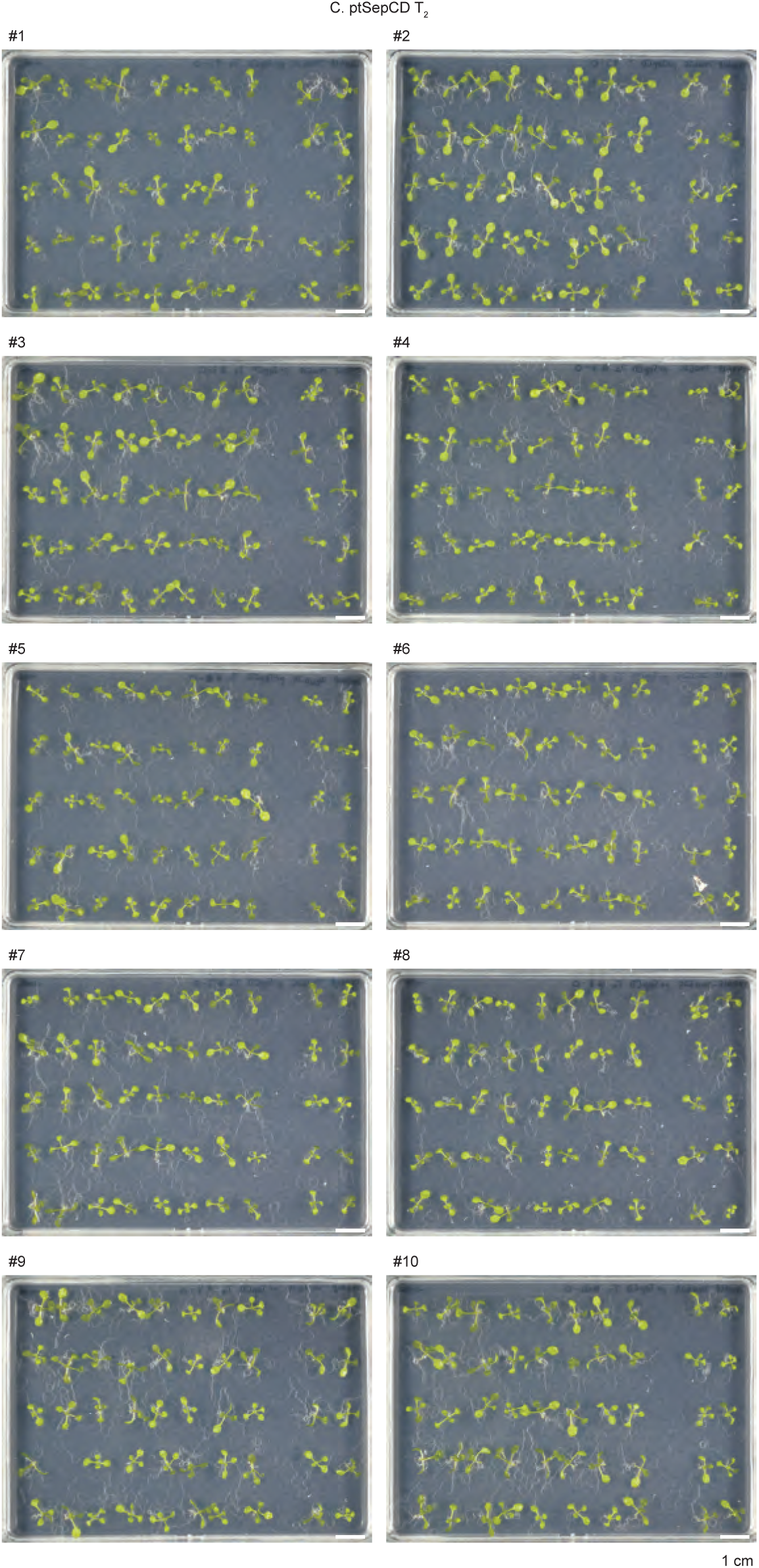
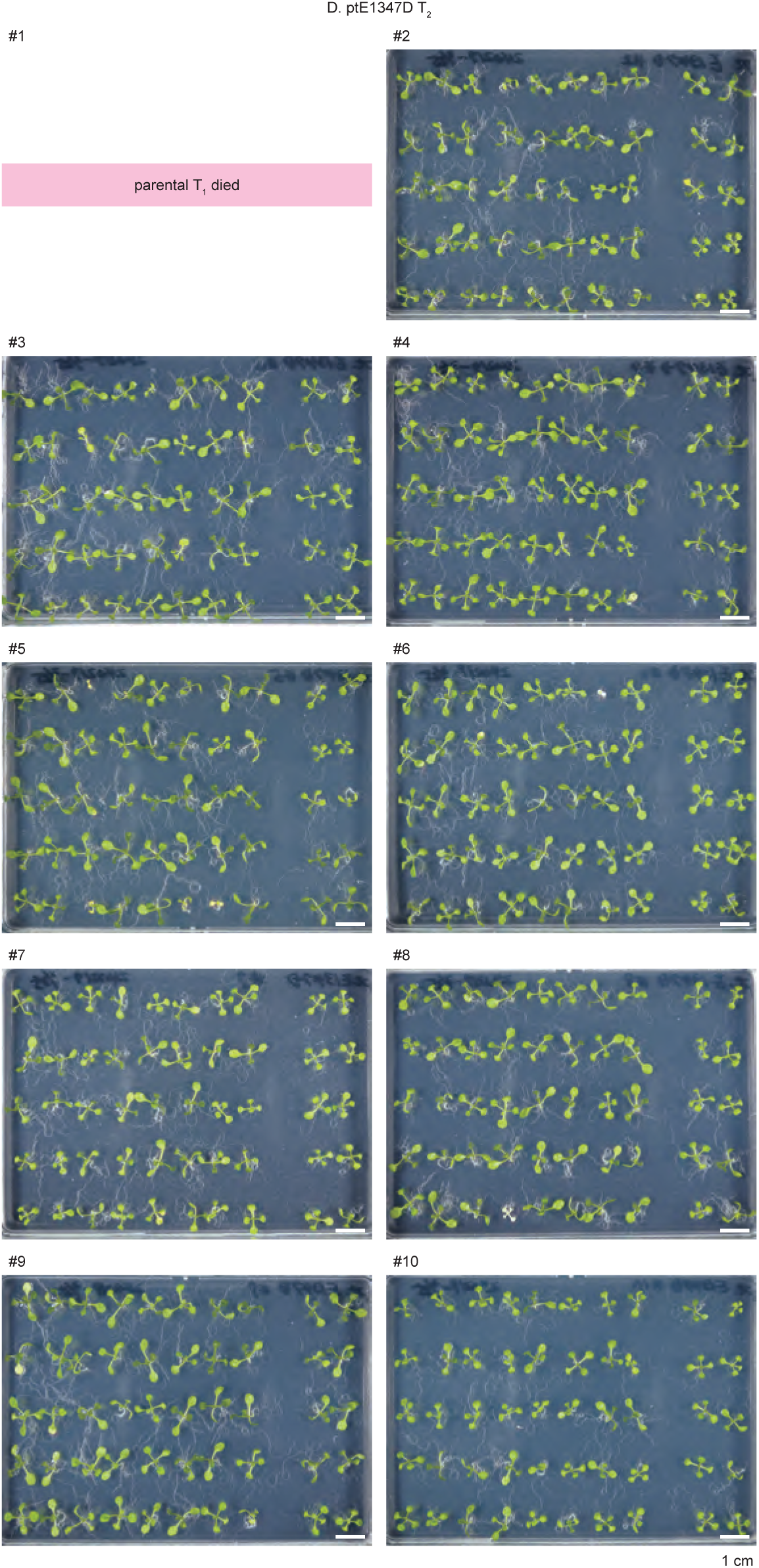
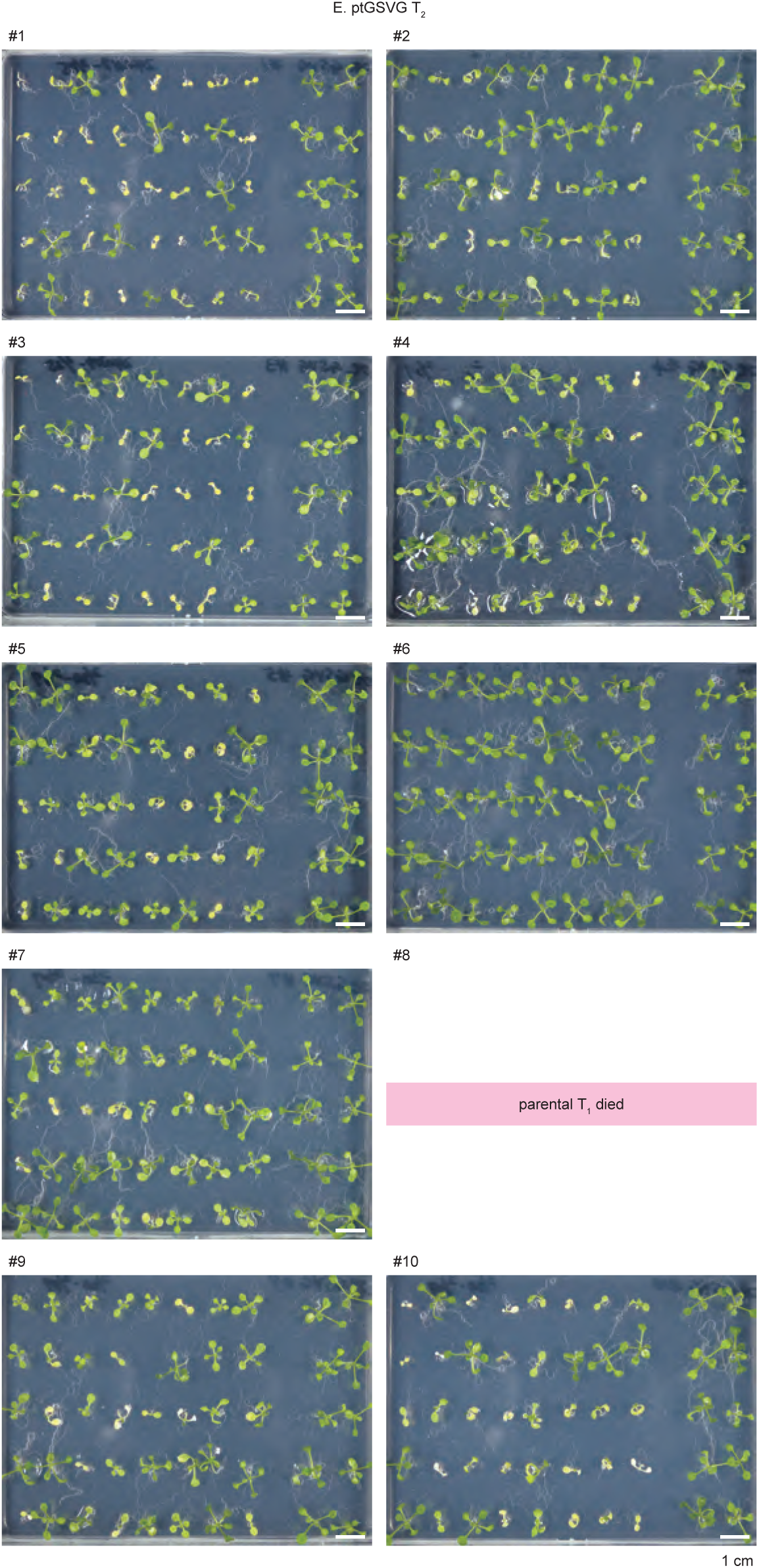
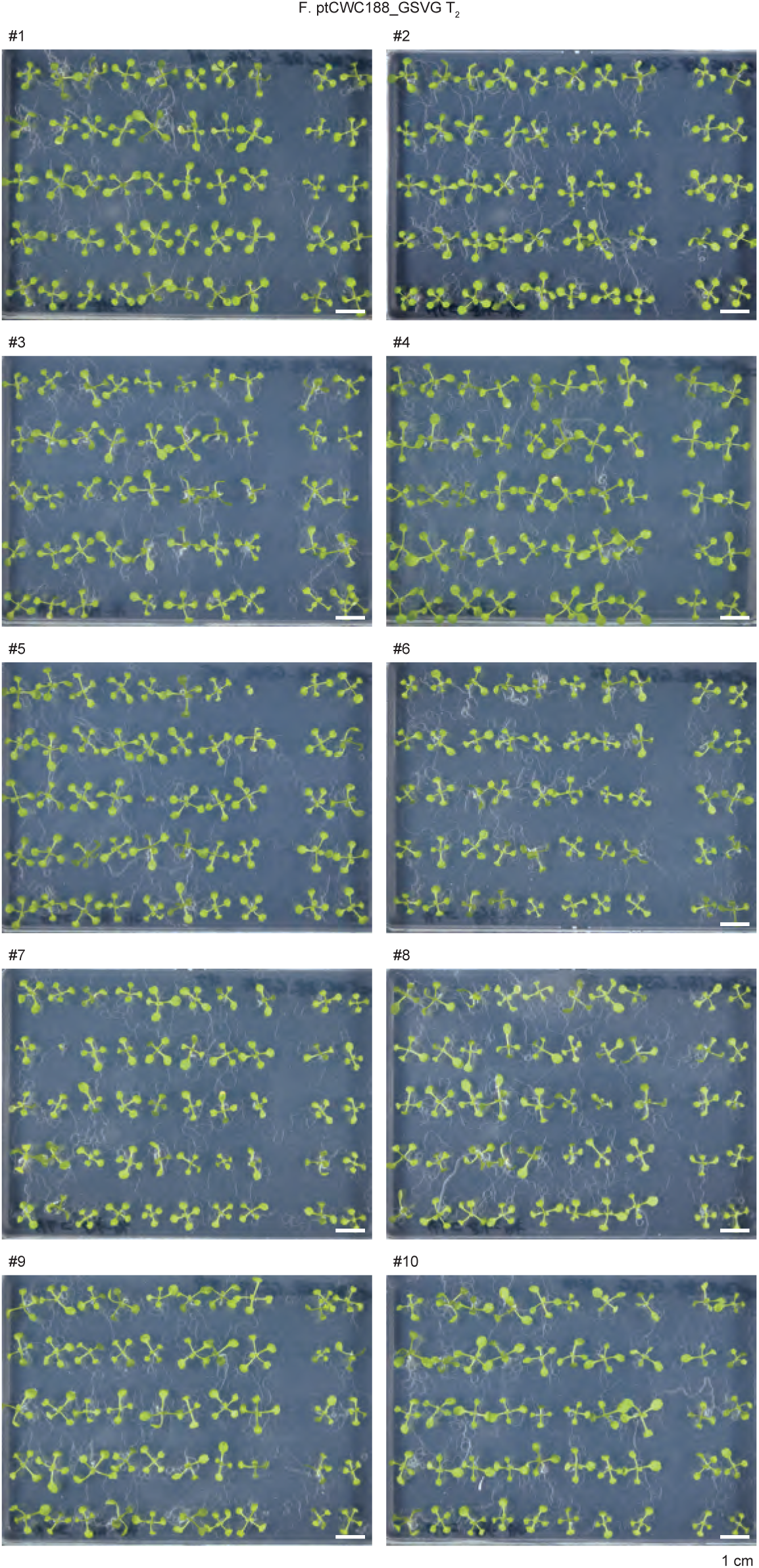
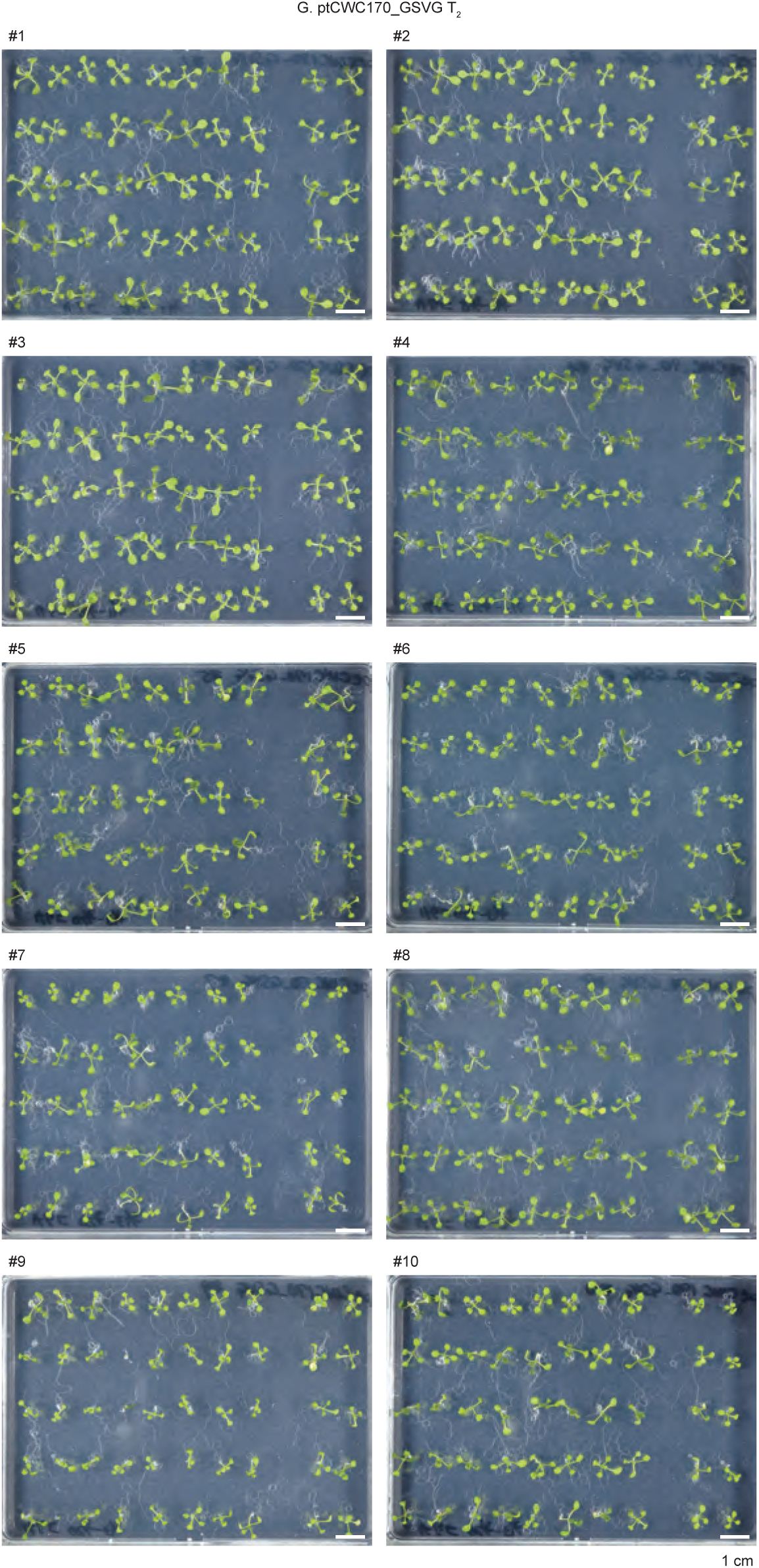
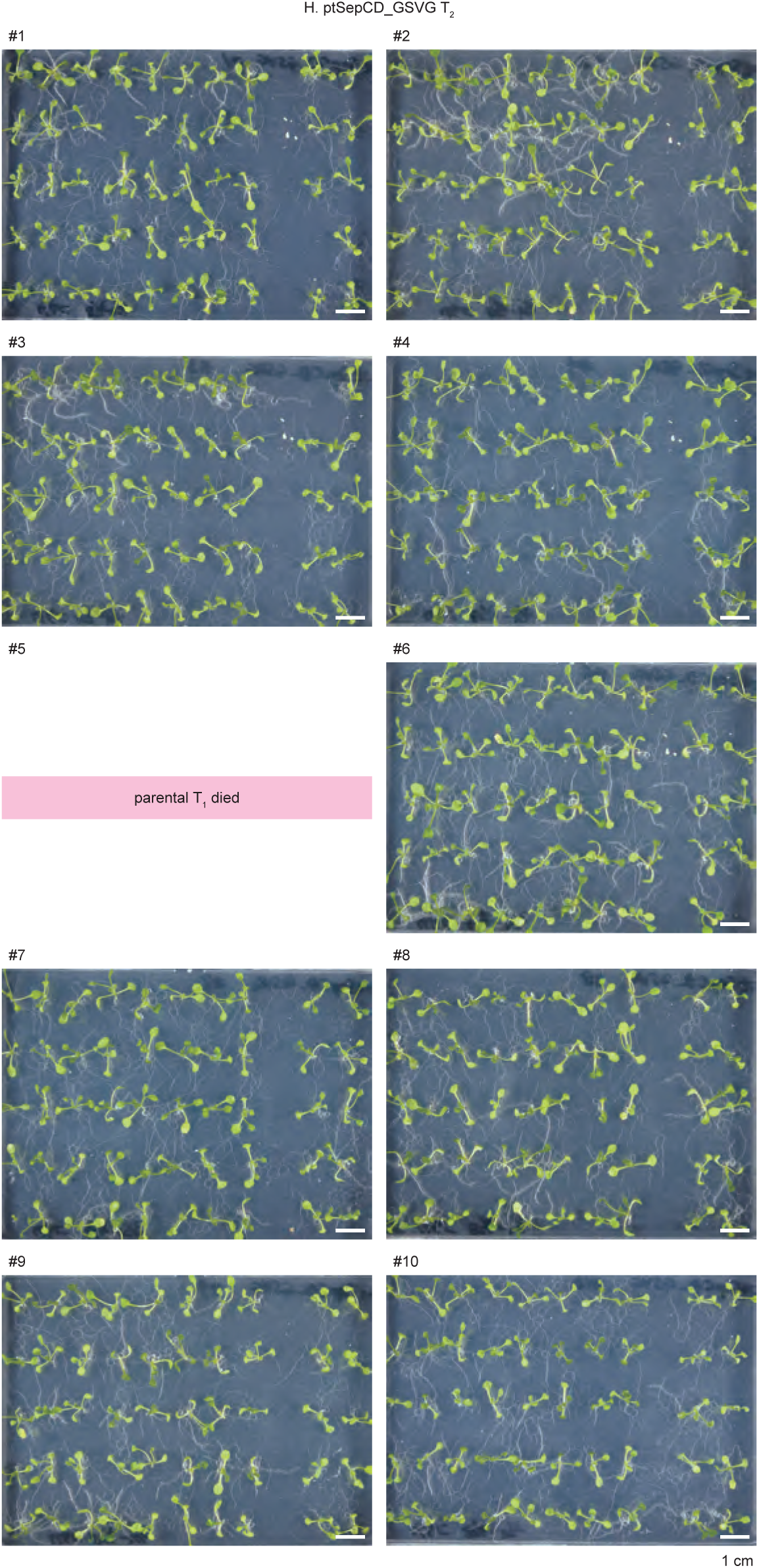
Photographs used for T_2_ visual observation phenotyping (supporting Table 1) 9 DAS. Magenta highlights indicate that the parental T_1_ died, subsequently unable to observe the T_2_ progeny. **A.** Legends. **B.** *msh1* mutant populations and Col-0. **C.** Mitochondria-mutagenized T_2_ populations. **D.** Plastids-mutagenized T_2_ populations.

**Supplementary Fig. S7.**
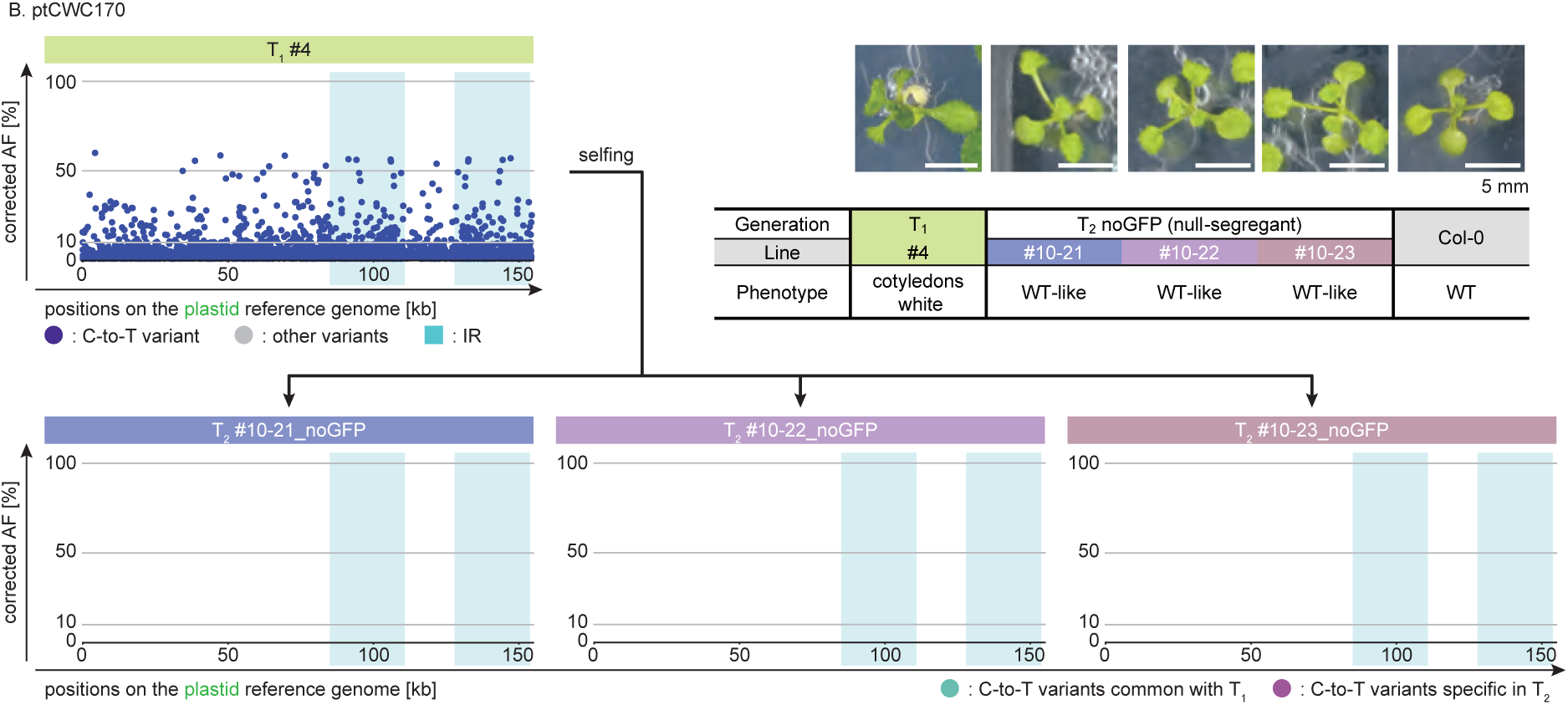
Whole genome NGS and SNP call analysis of three T_2_ individuals derived from B. ptCWC170 #4, and their phenotypes (supporting Fig. 5) Phenotypes are at 14 DAS. For more detailed sequencing information, see Supplementary Data Set 2.

**Supplementary Fig. S8.**
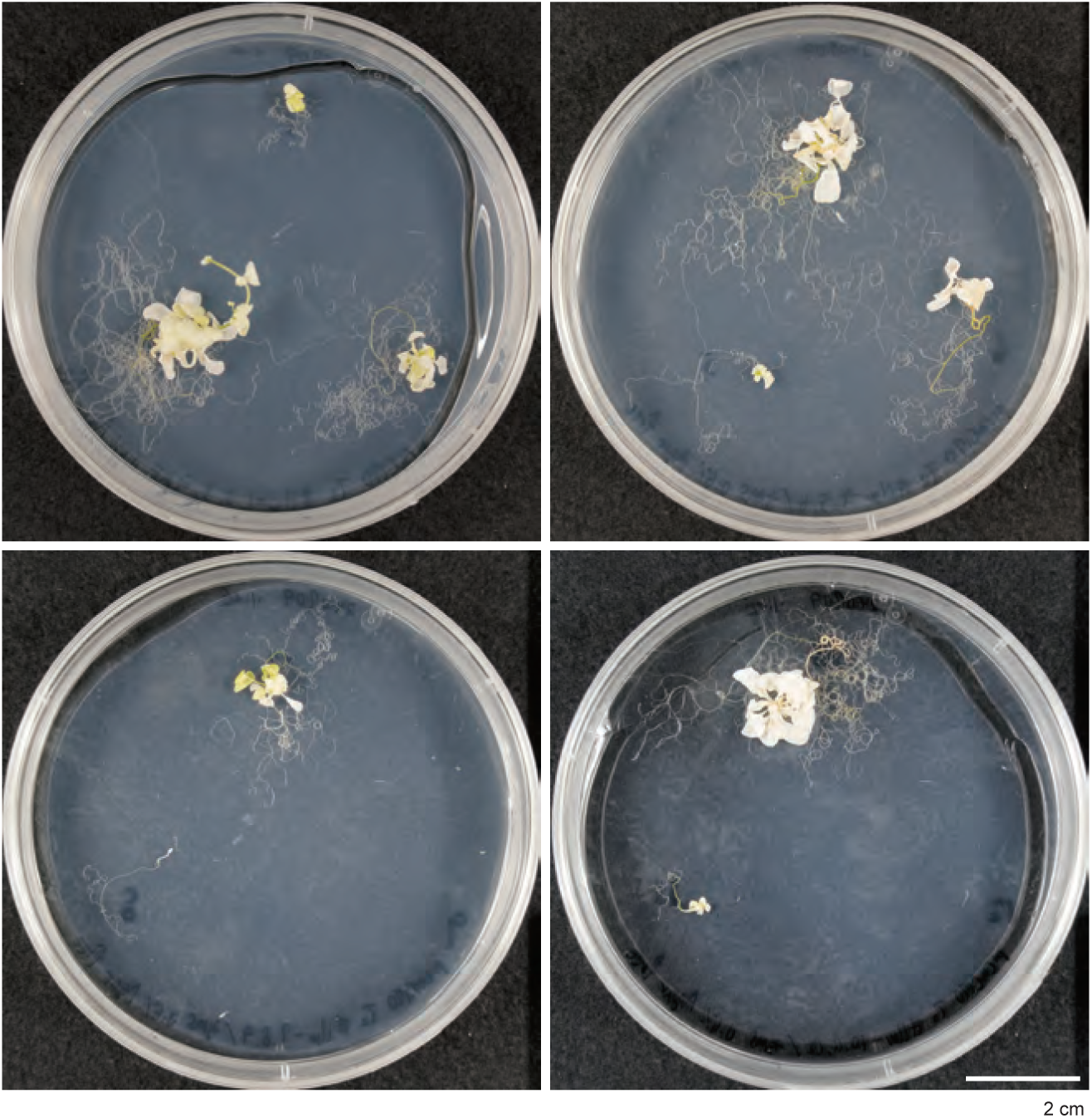
The phenotypes of T_2_ derived from white tissue, continued to be grown on 1/2 MS medium containing sucrose (supporting Fig. 6) 99 DAS. Col-0 (WT) is absent because it had already set seeds and died at 99 DAS.

**Supplementary Fig. S9.**
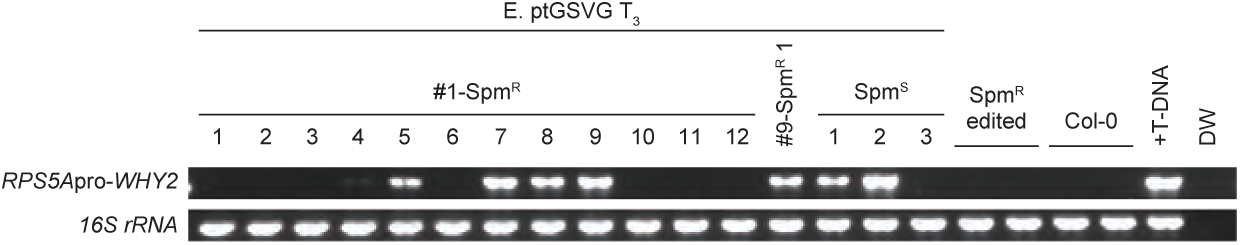
Gel images of bands of PCR products of T-DNA encoding WHY2-CD mutator, and *16S rRNA* (supporting Fig. 7) For information of the primers used, see Supplementary Table S8. DW, distilled water (negative control).

**Supplementary Fig. S10.**
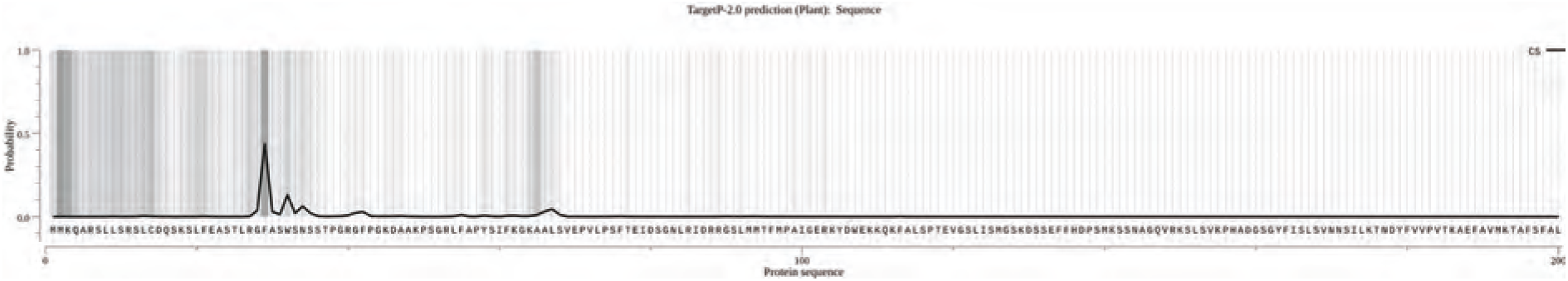
Prediction of WHY2 protein cleavage by TargetP 2.0 (supporting Materials & Methods) For constructs named “CWC,” cleavage is likely to occur at amino acids predicted to be cleaved by the organelle transfer signal sequence inherent in native WHY2 itself, in order to avoid the separation of WHY2-CD mutator peptides and the existence of split CDs as separate molecules. The query used the full-length amino acid sequence of WHY2 (Supplementary Sequences).

**Supplementary Fig. S11.**
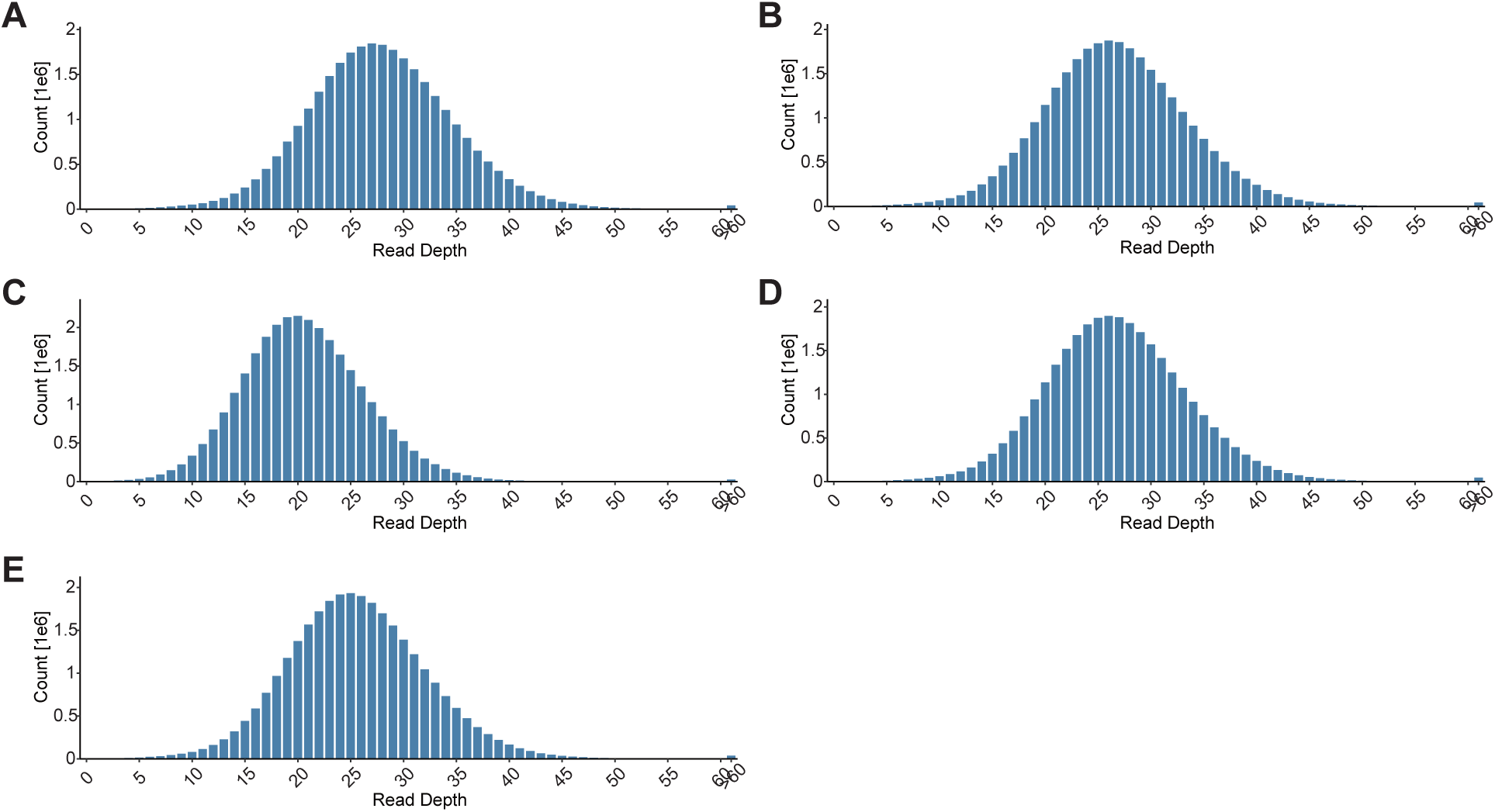
Read depth histograms on the chromosome 1 (supporting Materials & Methods) **A.** mtE1347D_bulk, **B.** ptE1347D_bulk, **C.** mtGSVG_bulk, **D.** ptGSVG_bulk, **E.** Col-0_bulk.

**Supplementary Fig. S12.**
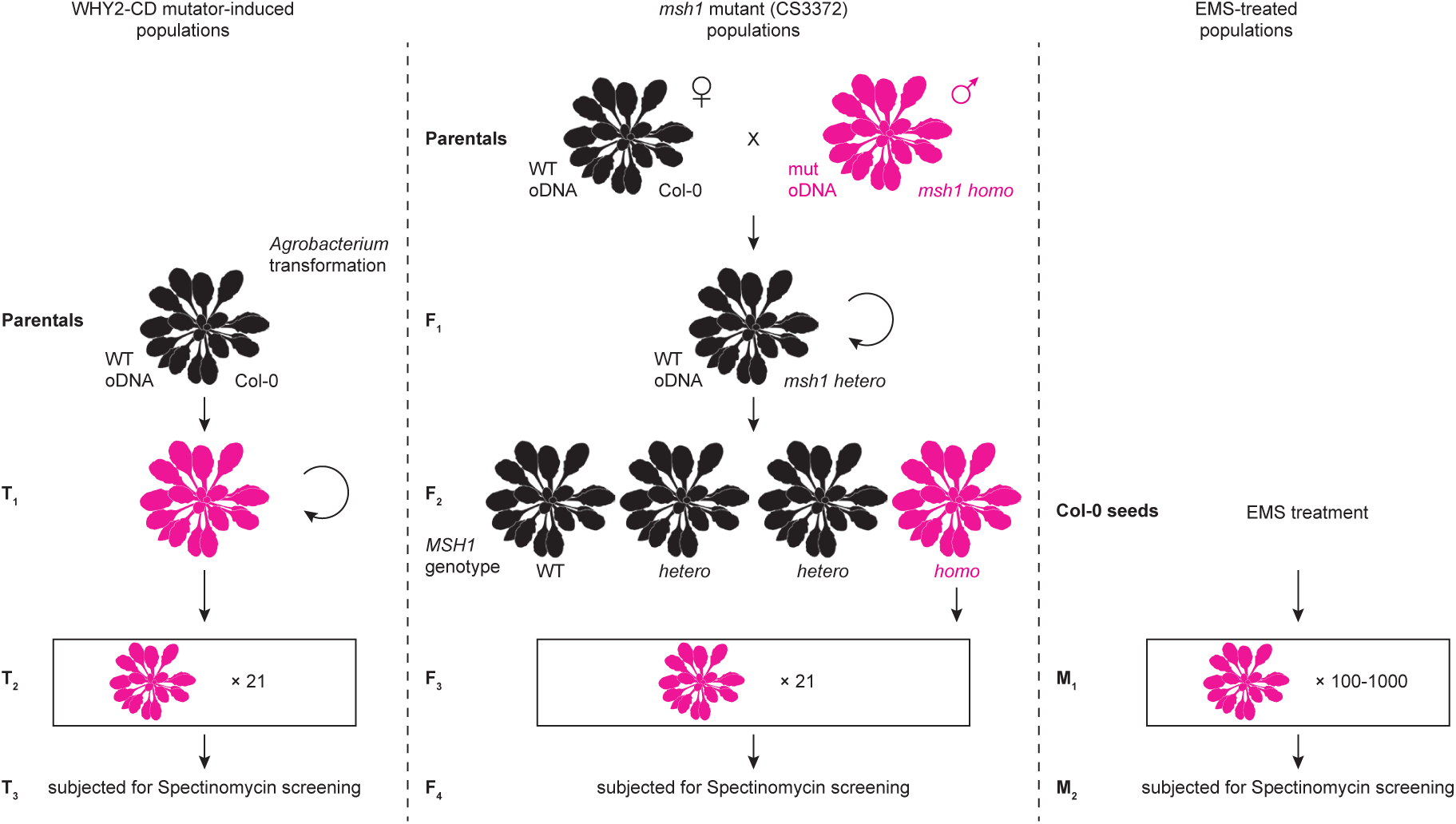
Method for preparing mutant populations used in spectinomycin screening (supporting Materials & Methods) The method shown in this figure was used to ensure that the number of generations in which mutations could accumulate in the organelle genome and the possibility of homoplasmic mutations occurring in the genome were consistent. Arabidopsis silhouette image is from PhyloPic (Mason McNair). This figure was partially modified from Wu *et al.*, 2020.

**Supplementary Table S1.**
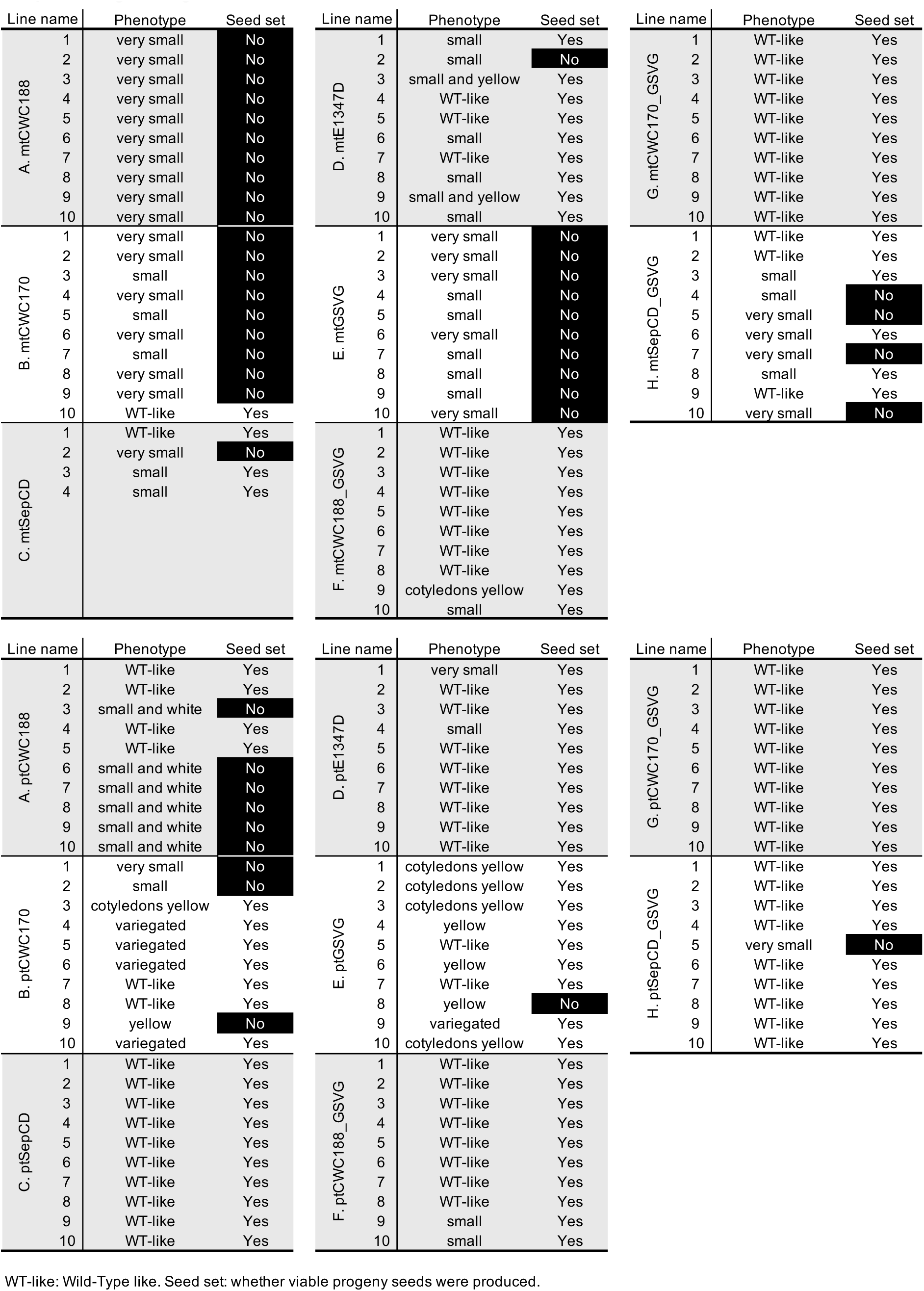
Phenotype description of T_1_ plants, subjected for Sanger sequencing in Fig. 2C, D.

**Supplementary Table S2.**
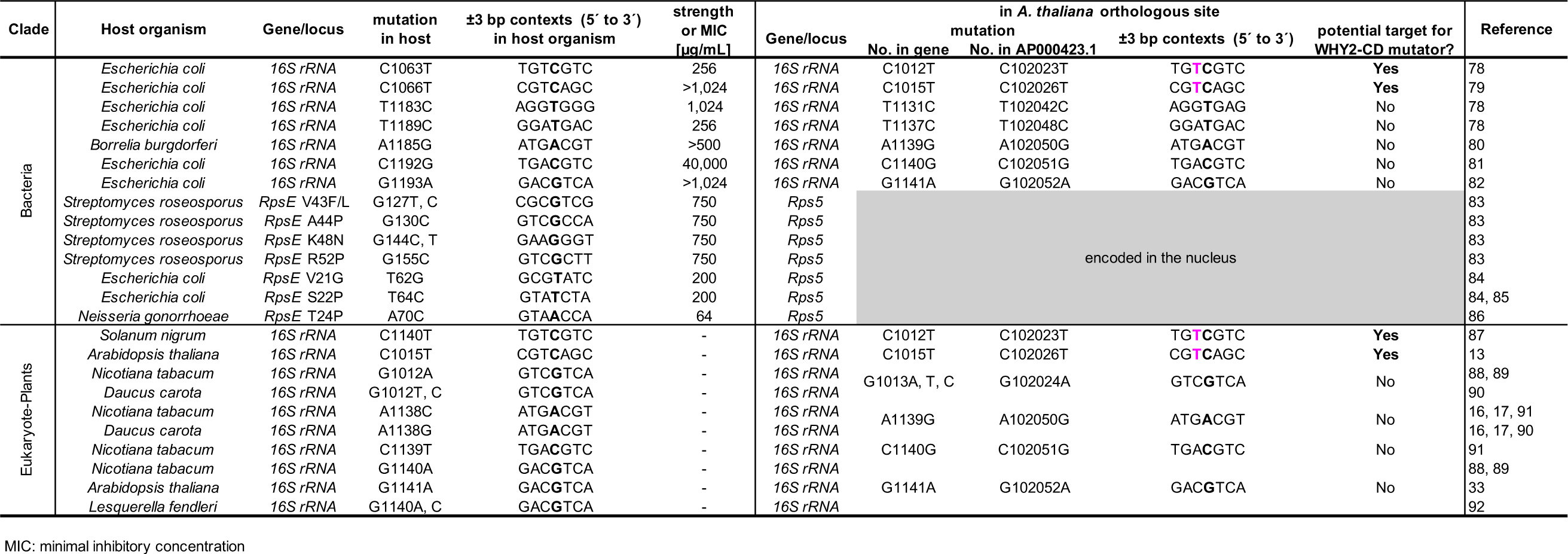
List of previously reported single nucleotide mutations conferring spectinomycin resistance.

**Supplementary Table S3.**
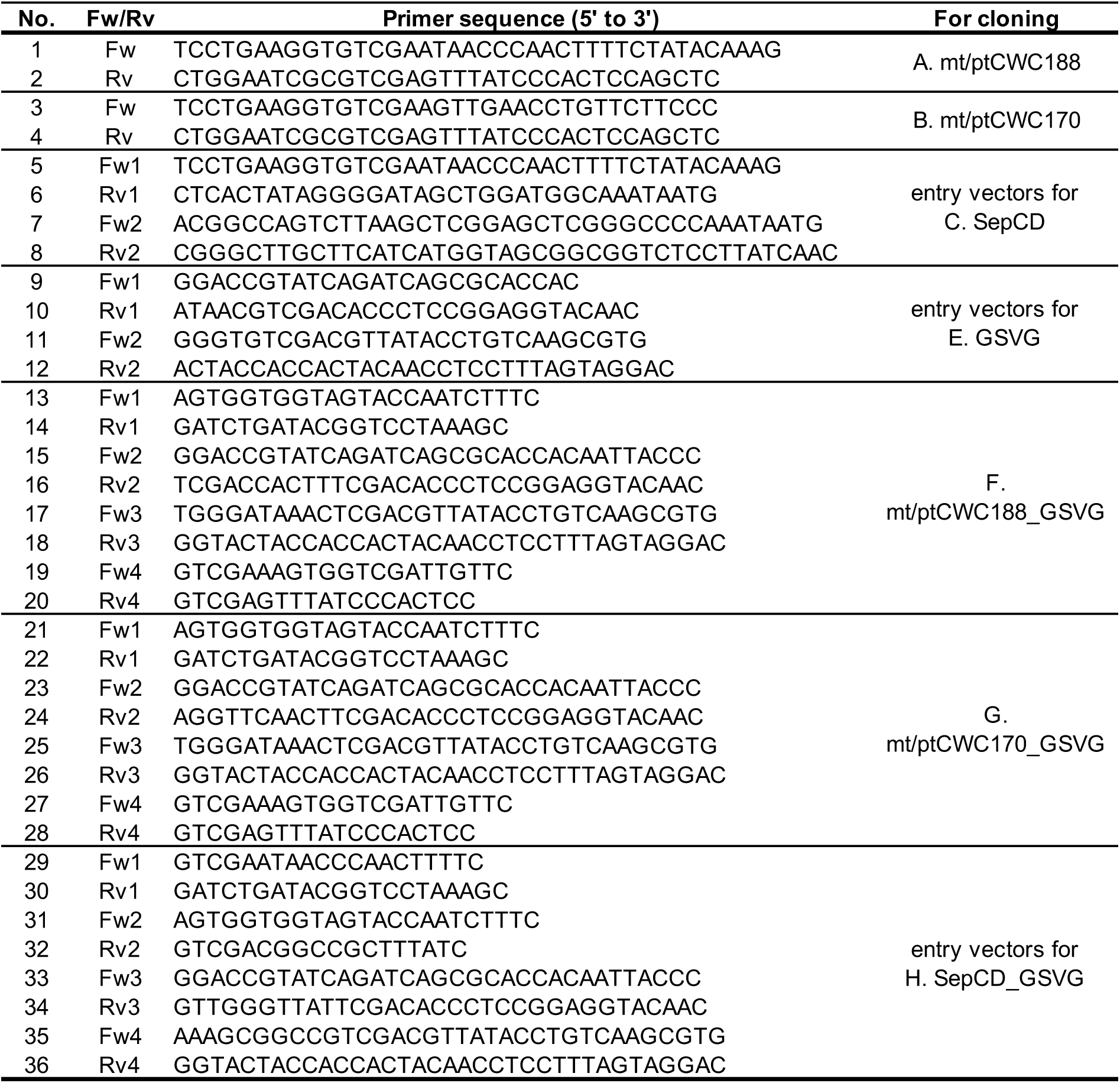
List of primers for vector constructions.

**Supplementary Table S4.**
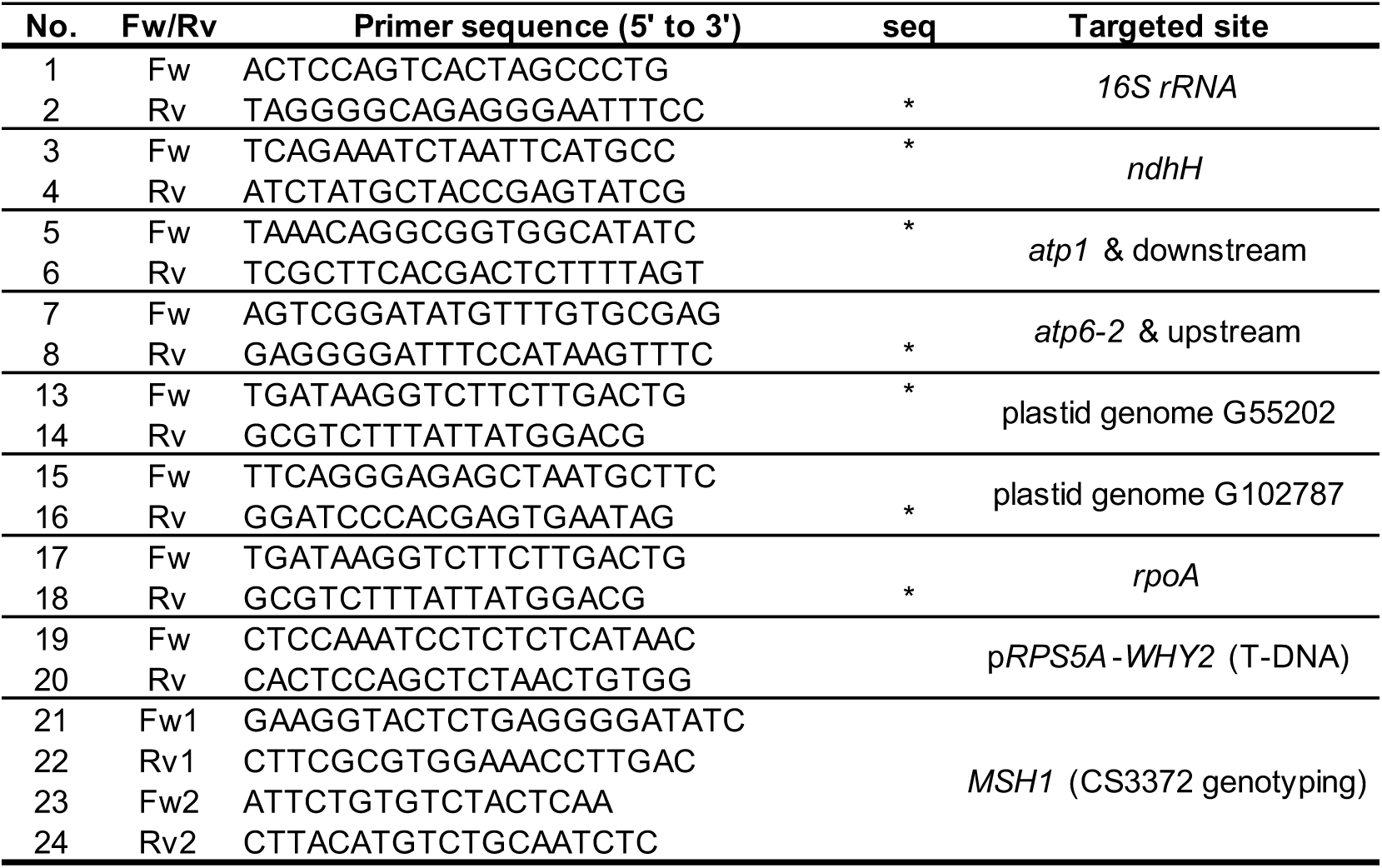
List of primers for Sanger sequencing & genotyping.

